# Sex differences in islet stress responses support female beta cell resilience

**DOI:** 10.1101/2022.05.10.491428

**Authors:** George P. Brownrigg, Yi Han Xia, Chieh Min Jamie Chu, Su Wang, Charlotte Chao, Jiashuo Aaron Zhang, Søs Skovsø, Evgeniy Panzhinskiy, Xiaoke Hu, James D. Johnson, Elizabeth J. Rideout

**Affiliations:** Department of Cellular and Physiological Sciences, Life Sciences Institute, The University of British Columbia, 2350 Health Sciences Mall, Vancouver, BC, V6T 1Z3, Canada

**Keywords:** pancreatic islets, β cells, diabetes mellitus, endoplasmic reticulum stress, protein synthesis, transcriptomics

## Abstract

**Objective:** Pancreatic β cells play a key role in glucose homeostasis; dysfunction of this critical cell type causes type 2 diabetes (T2D). Emerging evidence points to sex differences in β cells, but few studies have examined male-female differences in β cell stress responses and resilience across multiple contexts, including diabetes. Here, we address the need for high-quality information on sex differences in β cell/islet gene expression and function using both human and rodent samples.

**Methods:** We compared β cell gene expression and insulin secretion in donors living with T2D to non-diabetic donors in both males and females. In mice, we generated a well-powered islet RNAseq dataset from 20-week-old male and female siblings with equivalent insulin sensitivity. Because on our unbiased analysis of gene expression pointed to sex differences in endoplasmic reticulum (ER) stress response, we subjected islets isolated from age-matched male and female mice to thapsigargin treatment and monitored protein synthesis, cell death, and β cell insulin production and secretion. Transcriptomic and proteomic analyses were used to characterize sex differences in islet responses to ER stress.

**Results:** Our single-cell analysis of human β cells revealed sex-specific changes to gene expression and function in T2D, correlating with more robust insulin secretion in islets isolated from female donors living with T2D compared to male T2D donors. In mice, RNA sequencing revealed differential enrichment of unfolded protein response pathway-associated genes, where female islets showed higher expression of genes linked with protein synthesis, folding, and processing. This differential expression was biologically significant, as female islets were more resilient to ER stress induction with thapsigargin. Specifically, female islets maintained better insulin secretion and showed a distinct transcriptional response under ER stress compared with males.

**Conclusions:** Our data demonstrate that physiologically significant sex differences in β cell gene expression exist in both humans and mice, and that female β cells maintain better insulin production and secretion across multiple physiological and pathological contexts.

## 1. Introduction

Pancreatic β cells make and secrete insulin, an essential hormone required to maintain whole-body glucose homeostasis. Emerging evidence from multiple species suggests biological sex is an important, but often overlooked, factor that affects β cell biology (1–6). Large-scale surveys of gene expression in mice and humans show that differences exist between the sexes in the pancreas (7–9), in islets (10), and in β cells specifically (4,11). Humans also have a sex-specific β cell gene expression response to aging (12), and show male-female differences in pancreatic β cell number (6). With respect to β cell function, most data from rodent and human studies suggests glucose-stimulated insulin secretion is higher in females than in males (5,10,13–16). While male-female differences in peripheral insulin sensitivity (15,17–27) may contribute to these differences, sex-biased insulin secretion in humans persists in the context of equivalent insulin sensitivity between males and females (5). Whether sex differences in additional aspects of β cell gene expression and function similarly persist remains unclear, as insulin sensitivity is not routinely monitored across datasets showing sex differences in β cell biology.

Biological sex also affects the risk of developing T2D. Across many population groups, men are at a higher risk of developing T2D than women (28–31). Some of the differential risk is explained by lifestyle and cultural factors (31–33). Biological sex also plays a role, however, as the male-biased risk of developing diabetes-like phenotypes exists across multiple animal models (22,34–39). Despite a dominant role for β cell function in T2D pathogenesis (40,41), T2D- and stress-associated changes to β cell gene expression and function in each sex remain largely unexplored, as many studies on this topic did not include biological sex as a variable in their analysis (42–49). More detailed knowledge of β cell gene expression and function in physiological and pathological contexts is therefore a key first step toward understanding how sex differences in this important cell type may contribute to T2D risk.

The overall goal of our study was to provide detailed knowledge of β cell gene expression and function in both males and females across multiple contexts to advance our understanding of sex differences in this important cell type. Collectively, our data show significant sex differences in islet and β cell gene expression and stress responses in both humans and mice. These differences contribute to sex differences in β cell resilience, where we find female β cells maintain better insulin secretion in response to stress and T2D. Importantly, these differences cannot be fully explained by differential peripheral insulin sensitivity between the sexes, suggesting biological sex is an important variable to consider in studies on islet and β cell function.

## 2. Materials and methods

### 2.1. Animals

Mice were bred in-house or purchased from the Jackson Laboratory. Unless otherwise stated mouse islets were isolated from C57BL/6J mice aged 20-24 weeks. Animals were housed and studied in the UBC Modified Barrier Facility using protocols approved by the UBC Animal Care Committee and in accordance with international guidelines. Mice were housed on a 12-hour light/dark cycle with food and drinking water *ad libitum*. Mice were fed a regular chow diet (LabDiet #5053); 24.5% energy from protein, 13.1% energy from fat, and 62.4% energy from carbohydrates.

### 2.2. Islet Isolation, Culture, Dispersion and Treatment

Mouse islet isolations were performed by ductal collagenase injection followed by filtration and hand-picking, using modifications of the protocol described by Salvalaggio (50). Islets recovered overnight, in islet culture media (RPMI media with 11.1 mM D-glucose supplemented with 10% vol/vol fetal bovine serum (FBS) (Thermo: 12483020) and 1% vol/vol Penicillin-Streptomycin (P/S) (GIBCO: 15140-148)) at 37°C with 5% CO2. After four washes with Minimal Essential Medium [L-glutamine, calcium and magnesium free] (Corning: 15-015 CV) islets were dispersed with 0.01% trypsin and resuspended in islet culture media. Cell seedings were done as per the experimental procedure (protein synthesis: 20,000 cells per well, live cell imaging: 5,000 cells per well). ER stress was induced by treating islets with the SERCA inhibitor thapsigargin. For assays less than 24 hours, we used (11.1 mM D-glucose RPMI, 1% vol/vol P/S). For assays greater than 24 hours we used (11.1 mM D-glucose RPMI, 1% vol/vol P/S, 10% vol/vol FBS).

### 2.3. Analysis of protein synthesis

Dispersed islets were seeded into an optical 96-well plate (Perkin Elmer) at a density of approximately 20,000 cells per well islet culture media (11.1 mM D-glucose RPMI, 1% vol/vol P/S, 10% vol/vol FBS). 24 hours after seeding, treatments were applied in fresh islet culture media (11.1 mM D-glucose RPMI, 1% vol/vol P/S). After incubation, fresh culture media was applied (11.1 mM D-glucose RPMI, 1% vol/vol P/S), supplemented with 20 μM OPP (Invitrogen) and respective drug treatments. The assay was performed according to manufacturer’s instructions; cells were then imaged at 10x with an ImageXpress^MICRO^ high-content imager and analyzed with MetaXpress (Molecular Devices) to quantify the integrated staining intensity of OPP-Alexa Fluor 594 in cells identified by NuclearMask Blue Stain.

### 2.4. Live cell imaging

Dispersed islets were seeded into 384-well plates (Perkin Elmer) at a density of approximately 5,000 cells per well and allowed to adhere for 48 hours in islet culture media (11.1 mM D-glucose RPMI, 1% vol/vol P/S, 10% vol/vol FBS). Cells were stained with Hoechst 33342 (Sigma-Aldrich) (0.05 μg/mL) and propidium iodide (Sigma-Aldrich) (0.5 μg/mL) for one hour in islet culture media (11.1 mM D-glucose RPMI, 1% vol/vol P/S, 10% vol/vol FBS) prior to the addition of treatments and imaging. 384-well plates were placed into environmentally controlled (37°C, 5% CO2) ImageXpress^MICRO^ high content imaging system. To measure cell death, islet cells were imaged every 2 hours for 84 hours, and MetaXpress software was used to quantify cell death, defined as the number of Propidium Iodide-positive/Hoechst 33342-positive cells. To measure *Ins2* gene activity, dispersed islets from *Ins2*^GFP/WT^ mice aged 21-23 weeks were used (51). Islet cells were imaged every 30 minutes for 60 hours. MetaXpress analysis software and custom R scripts were used to perform single-cell tracking of *Ins2*^GFP/WT^ β cells as previously described (51).

### 2.5. Western blot

After a 24-hour treatment with 1 μM Tg in islet culture media (11.1 mM D-glucose RPMI, 1% vol/vol P/S, 10% vol/vol FBS), mouse islets were sonicated in RIPA lysis buffer (150 mM NaCl, 1% Nonidet P-40, 0.5% DOC, 0.1% SDS, 50 mM Tris (pH 7.4), 2 mM EGTA, 2 mM Na_3_VO_4_, and 2 mM NaF supplemented with complete mini protease inhibitor cocktail (Roche, Laval, QC)). Protein lysates were incubated in Laemmli loading buffer (Thermo, J61337AC) at 95°C for 5 minutes and resolved by SDS-PAGE. Proteins were then transferred to PVDF membranes (BioRad, CA) and probed with antibodies against HSPA5 (1:1000, Cat. #3183, Cell Signalling), eIF2α (1:1000, Cat. #2103, Cell Signalling), phospho-eIF2α (1:1000, Cat. #3398, Cell Signalling), IRE1α (1:1000, Cat. #3294, Cell Signalling), phospho-IRE1α (1:1000, Cat. #PA1-16927, Thermo Fisher Scientific), CHOP (1:1000, #ab11419, Abcam), β-actin (1:1000, NB600-501, Novus Biologicals). The signals were detected by secondary HRP-conjugated antibodies (Anti-mouse, Cat. #7076; Anti-rabbit, Cat. #7074; CST) and either Pierce ECL Western Blotting Substrate (Thermo Fisher Scientific) or Forte (Immobilon). Protein band intensities were quantified using Image Studio (LI-COR).

### 2.6. Islet Secretion and Content

Glucose-stimulated insulin/proinsulin production and secretion were assessed using size-matched islets (five islets per well, in triplicate) seeded into 96-well V-bottom Tissue Culture Treated Microplates (Corning: #CLS3894). Islets were allowed to adhere for 48 hours in culture media (11.1 mM D-glucose RPMI, 1% vol/vol P/S, 10% vol/vol FBS). Islets were washed with Krebs-Ringer Buffer (KRB; 129 mM NaCl, 4.8 mM KCl, 1.2 mM MgSO_4_, 1.2 mM KH_2_PO_4_, 2.5 mM CaCl_2_, 5 mM NaHCO3, 10 mM HEPES, 0.5% bovine serum albumin) containing 3 mM glucose then pre-incubated for 4 hours in 3 mM glucose KRB. 1 μM Tg was added to the 3 mM low glucose pre-incubation buffer 4 hours prior, 2 hours prior, or at the start of the low glucose incubation period. Islets were incubated in KRB with 3 mM glucose then 20 mM glucose for 45 minutes each. Supernatant was collected after each stimulation. Islet insulin and proinsulin content was extracted by freeze-thawing in 100 μL of acid ethanol, then the plates were shaken at 1200 rpm for 10 minutes at 4°C to lyse the islets. Insulin was measured by Rodent Insulin Chemiluminescent ELISA (ALPCO: 80-INSMR) and proinsulin by Rat/Mouse Proinsulin ELISA (Mercodia: 10-1232-01). Measurements were performed on a Spark plate reader (TECAN).

### 2.7. Blood collection and in vivo analysis of glucose homeostasis and insulin secretion

Mice were fasted for 6 hours prior to glucose and insulin tolerance tests. During glucose and insulin tolerance tests, tail blood was collected for blood glucose measurements using a glucometer (One Touch Ultra 2 Glucometer, Lifescan, Canada). For intraperitoneal (i.p.) glucose tolerance tests, the glucose dose was 2 g glucose/kg of body mass. For insulin tolerance tests, the insulin dose was 0.75U insulin/kg body mass. For measurements of *in vivo* glucose-stimulated insulin secretion, femoral blood was collected after i.p. injection of 2 g glucose/kg body mass. Blood samples were kept on ice during collection, centrifuged at 2000 rpm for 10 minutes at 4°C and stored as plasma at −20°C. Plasma samples were analysed for insulin using Rodent Insulin Chemiluminescent ELISA (ALPCO: 80-INSMR).

### 2.8. RNA sequencing

To assess basal transcriptional differences islets from male and female mice (n=9, 8) were snap frozen and stored at −80°C until RNA extraction. To assess Tg-induced transcriptional changes islets from each mouse were treated with DMSO or Tg for 6- or 12-hours in culture media (11.1 mM D-glucose RPMI, 1% vol/vol P/S) (8 groups, n=3-4 per group, each n represents pooled islet RNA from two mice). Islets were frozen at −80°C in 100 μL of RLT buffer (Qiagen) with beta mercaptoethanol (1%). RNA was isolated using RNeasy Mini Kit (Qiagen #74106) according to manufacturer’s instructions. RNA sequencing was performed at the UBC Biomedical Research Centre Sequencing Core. Sample quality control was performed using the Agilent 2100 Bioanalyzer System (RNA Pico LabChip Kit). Qualifying samples were prepped following the standard protocol for the NEBNext Ultra II Stranded mRNA (New England Biolabs). Sequencing was performed on the Illumina NextSeq 500 with Paired End 42bp × 42bp reads. Demultiplexed read sequences were then aligned to the reference sequence (UCSC mm10) using STAR aligner (v 2.5.0b) (52). Gene differential expression was analyzed using DESeq2 R package (53). Pathway enrichment analysis were performed using Reactome (54). Over-representation analysis was performed using NetworkAnalyst3.0 (www.networkanalyst.ca) (55).

### 2.9. Proteomics

Islets were treated with DMSO or Tg for 6 hours in islet culture media (11.1 mM D-glucose RPMI, 1% vol/vol P/S) (4 groups, n=5-7 per group, each n represents pooled islets from two mice). Islet pellets were frozen at −80°C in 100 μL of SDS lysis buffer (4% SDS, 100 mM Tris, pH 8) and the proteins in each sample were precipitated using acetone. University of Victoria proteomics service performed non-targeted quantitative proteomic analysis using data-independent acquisition (DIA) with LC-MS/MS on an Orbitrap mass spectrometer. A mouse FASTA database was downloaded from Uniprot (http://uniprot.org). This file was used with the 6 gas phase fraction files from the analysis of the chromatogram library sample to create a mouse islet specific chromatogram library using the EncyclopeDIA (v 1.2.2) software package (Searle et al, 2018). This chromatogram library file was then used to perform identification and quantitation of the proteins in the samples again using EncyclopeDIA with Overlapping DIA as the acquisition type, trypsin used as the enzyme, CID/HCD as the fragmentation, 10 ppm mass tolerances for the precursor, fragment, and library mass tolerances. The Percolator version used was 3.10. The precursor FDR rate was set to 1%. Protein abundances were log2 transformed, imputation was performed for missing values, then proteins were normalized to median sample intensities. Gene differential expression was analyzed using limma in Perseus (56).

### 2.10. Data from HPAP

To compare sex differences in dynamic insulin secretion, data acquired was from the Human Pancreas Analysis Program (HPAP-RRID:SCR_016202) Database (https://hpap.pmacs.upenn.edu), a Human Islet Research Network (RRID:SCR_014393) consortium (UC4-DK-112217, U01-DK-123594, UC4-DK-112232, and U01-DK-123716).

### 2.11. Statistical Analysis

Statistical analyses and data presentation were carried out using GraphPad Prism 9 (GraphPad Software, San Diego, CA, USA) or R (v 4.1.1). Student’s *t*-tests or two-way ANOVAs were used for parametric data. A Mann-Whitney test was used for non-parametric data. Statistical tests are indicated in the figure legends. For all statistical analyses, differences were considered significant if the p-value was less than 0.05. *: p<0.05; ** p<0.01; *** p<0.001. Data were presented as means ± SEM with individual data points from biological replicates.

## 3. Results

### 3.1. Sex differences in β cell transcriptional and functional responses in ND and T2D human islets

To define β cell-specific gene expression changes in T2D in each sex, we used a recently compiled meta-analysis of publicly available scRNAseq datasets from male and female human islets (57). In line with prior reports (12), non-diabetic (ND) and T2D β cells showed significant transcriptional differences. In β cells isolated from female T2D donors, mRNA levels of 127 genes were significantly different from ND female donors (77 downregulated, 50 upregulated in T2D) (Figure 1A-C). In β cells isolated from male T2D donors, 462 genes were differentially expressed compared with male ND donors (138 downregulated, 324 upregulated in T2D) (Figure 1A-C). Of the 660 genes that were differentially regulated in T2D, 71 were differentially regulated in both males and females (15 downregulated, 56 upregulated in T2D) (Figure 1A-C); however, the fold change for these 71 shared genes was different between males and females (Figure S1A; Supplementary file 1). This suggests that for shared genes, the magnitude of gene expression changes in T2D was not the same between the sexes. Beyond shared genes, we observed that the majority of differentially expressed genes in T2D (589/660) were unique to either males or females (Figure S1B, C; Supplementary file 1). Given that the most prominent gene expression changes in T2D were found in genes that were unique to one sex (Figure S2A, B; Supplementary file 1), these data suggest there are important sex differences in the β cell gene expression response to T2D.

**Figure 1.**
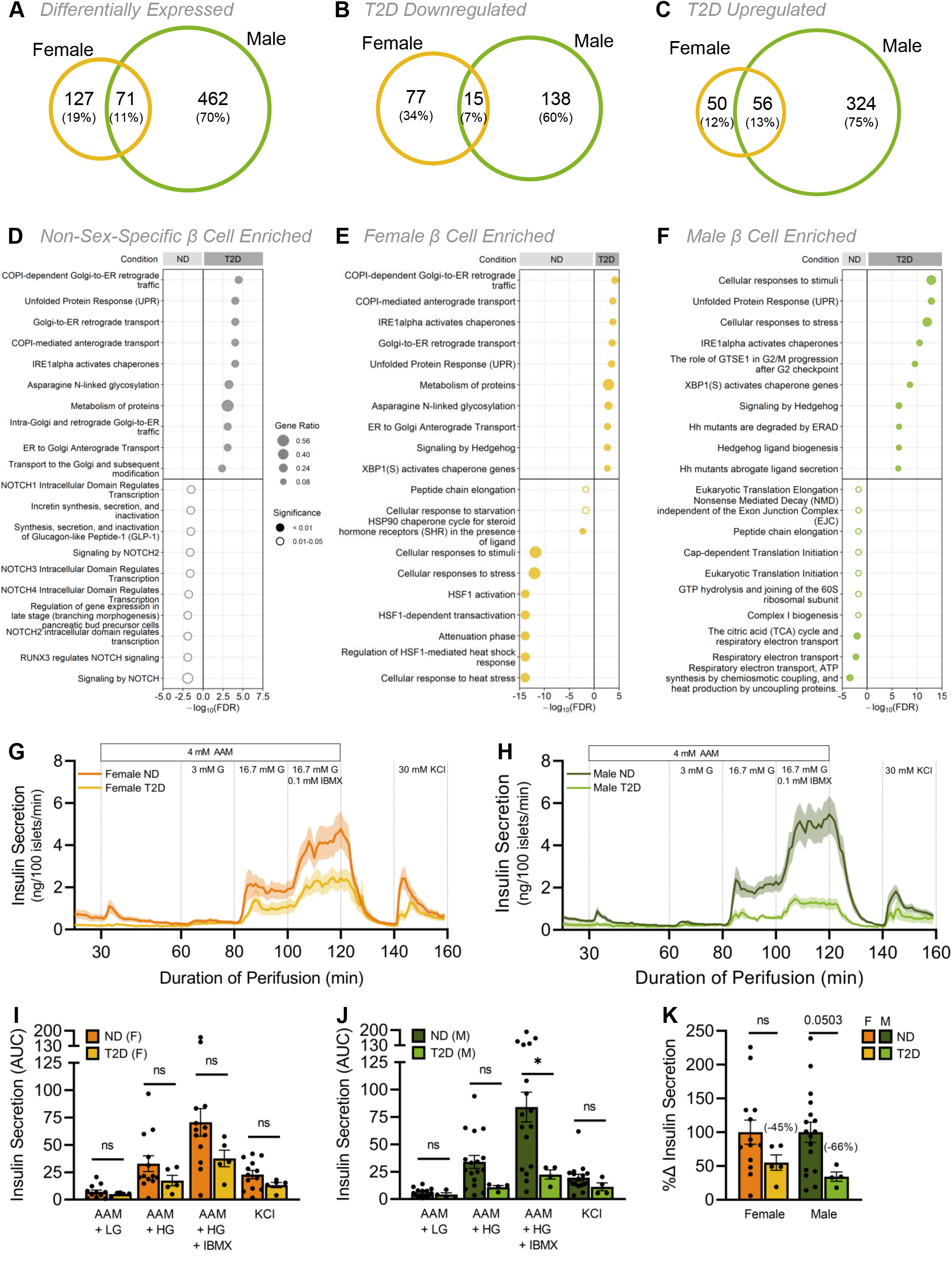
Sex differences in human islet transcriptomic and functional responses in type 2 diabetes. scRNAseq data from male and female human β cells. For donor metadata see Supplementary file 6. (A-C) Venn diagrams compare the number of significantly differentially expressed genes between ND and T2D donors (*p*-adj<0.05). All differentially expressed genes (A), downregulated genes (B), upregulated genes (C) in T2D human β cells. For complete gene lists see Supplementary file 1. (D-F) Top 10 significantly enriched Reactome pathways (ND vs T2D) from non-sex-specific (D), female (E), or male (F) significantly differentially expressed genes (*p*-adj< 0.05). Gene ratio is calculated as *k/n*, where *k* is the number of genes identified in each Reactome pathway, and *n* is the number of genes from the submitted gene list participating in any Reactome pathway. For complete Reactome pathway lists see Supplementary file 1. (G-K) Human islet perifusion data from the Human Pancreas Analysis Program in ND and T2D donor islets in females (F, I) and males (G, H). 3 mM glucose (3 mM G); 16.7 mM glucose (16.7 mM G); 0.1 mM isobutylmethylxanthine (0.1 mM IBMX); 30 mM potassium chloride (30 mM KCl); 4 mM amino acid mixture (4 mM AAM; mM: 0.44 alanine, 0.19 arginine, 0.038 aspartate, 0.094 citrulline, 0.12 glutamate, 0.30 glycine, 0.077 histidine, 0.094 isoleucine, 0.16 leucine, 0.37 lysine, 0.05 methionine, 0.70 ornithine, 0.08 phenylalanine, 0.35 proline, 0.57 serine, 0.27 threonine, 0.073 tryptophan, and 0.20 valine, 2 mM glutamine). (I-K) Quantification of area under the curve (AUC) is shown for the various stimulatory media in females (I), males (J) and donors with T2D (K). (I) In females, insulin secretion from ND islets was not significantly higher than T2D islets. (J) In males, insulin secretion from ND islets was significantly higher than T2D islets under 4 mM AAM +16.7 mM glucose (HG) + 0.1 mM IBMX stimulation (p=0.0442; unpaired Student’s *t*-test). (K) Total insulin secretion was lower in T2D male islets than ND male islets (p=0.0503; unpaired Student’s *t*-test). * indicates p<0.05; ns indicates not significant; error bars indicate SEM.

To determine which biological pathways were altered in β cells of T2D donors in each sex, we performed pathway enrichment analysis. Genes that were upregulated in β cells isolated from T2D donors included genes involved in Golgi-ER transport and the unfolded protein response (UPR) pathways (Figure 1D-F; Supplementary file 1). While these biological pathways were significantly upregulated in T2D in both males and females, ~75% of the differentially regulated genes in these categories were unique to each sex (Table 1). Genes that were downregulated in β cells from T2D donors revealed further differences between the sexes: biological pathways downregulated in β cells from female T2D donors included cellular responses to stress and to stimuli (Figure 1E; Supplementary file 1), whereas β cells from male T2D donors showed downregulation of pathways associated with respiratory electron transport and translation initiation (Figure 1F; Supplementary file 1). Thus, our analysis suggests that sex-biased β cell gene expression responses to T2D may influence different cellular processes in males and females.

**Table 1.** Human β cell pathway gene numbers. The number of genes corresponding to each T2D upregulated pathway in males, females or both sexes.

| Pathway Name | Number of Pathway Genes |  |  |
| --- | --- | --- | --- |
|  | Unique Male | Common | Unique Female |
| Asparagine N-linked glycosylation | 16 | 9 | 4 |
| Cellular responses to stimuli | 49 | 9 | 6 |
| Cellular responses to stress | 47 | 9 | 6 |
| COPI-dependent Golgi-to-ER retrograde traffic | 7 | 6 | 2 |
| COPI-mediated anterograde transport | 7 | 5 | 2 |
| ER to Golgi Anterograde Transport | 10 | 5 | 2 |
| Golgi-to-ER retrograde transport | 7 | 6 | 2 |
| Hedgehog ligand biogenesis | 12 | 4 | 1 |
| Hh mutants abrogate ligand secretion | 12 | 3 | 1 |
| Hh mutants are degraded by ERAD | 12 | 3 | 1 |
| IRE1alpha activates chaperones | 9 | 3 | 1 |
| Metabolism of proteins | 69 | 20 | 13 |
| Signaling by Hedgehog | 16 | 6 | 2 |
| The role of GTSE1 in G2/M progression after G2 checkpoint | 14 | 4 | 1 |
| Unfolded Protein Response (UPR) | 13 | 3 | 1 |
| XBP1(S) activates chaperone genes | 8 | 3 | 1 |

The sex-biased β cell transcriptional response in T2D prompted us to compare glucose-stimulated insulin secretion in each sex from ND and T2D human islets using data from the Human Pancreas Analysis Program database (58). In ND donors, islets from males and females showed similar patterns of insulin secretion in response to various stimulatory media (Figure 1G, H). In donors living with T2D, we found that insulin secretion was impaired to a greater degree in islets from males than in females (Figure 1G-K). Indeed, in male but not female islets, insulin secretion was significantly lower in donors with T2D following stimulation with both high glucose and IBMX (Figure 1I, J), which potentiates insulin secretion by increasing cAMP levels to a similar degree as the incretins (59). Human islets from female donors living with T2D therefore show better β insulin release than islets from males living with T2D (Figure 1K). Indeed, while diabetes status was the main donor characteristic that correlated with changes in insulin secretion (Figure S3A), we noted that in T2D sex and age were two donor characteristics showing trends toward an effect on insulin secretion (Figure S3B). Combined with our β cell gene expression data, these findings suggest that β cell transcriptional and functional responses in T2D are not shared between the sexes.

### 3.2. Sex differences in UPR-associated gene expression in mouse islets

Our unbiased analysis of human β cell gene expression and function in T2D revealed differences between male and female donors living with T2D. Because human β cell gene expression and function can be affected by factors such as peripheral insulin sensitivity, disease processes, and medication (31,33), we investigated sex differences in β cell gene expression and function in another context. We generated a well-powered islet RNAseq dataset from 20-week-old male and female C57BL/6J mice, an age where we show insulin sensitivity is equivalent between the sexes (Figure S4A). Principal component analysis and unsupervised clustering clearly separated male and female islets on the basis of gene expression (Figure 2A; Figure S5A). We found that 17.7% (3268/18938) of genes were differentially expressed between the sexes (1648 upregulated in females, 1620 upregulated in males), in line with estimates of sex-biased gene expression in other tissues (60,61). Overrepresentation and pathway enrichment analysis both identified UPR-associated pathways as a biological process that differed significantly between the sexes, where the majority of genes in this category were enriched in female islets (Figure 2B, C; Supplementary file 2). Additional genes that were enriched in female islets were those associated with the gene ontology term “Cellular response to ER stress” (GO:0034976), which included many genes involved in regulating protein synthesis (Figure 2D). For example, females showed significantly higher levels of most ribosomal protein genes (Figure 2E). Further genes enriched in females included those associated with protein folding, protein processing, and quality control (Figure 2D). Given that protein synthesis, processing, and folding capacity are intrinsically important for multiple islet cell types (62–65), including β cells (66,67), this suggests female islets may have a larger protein production and folding capacity than male islets.

**Figure 2.**
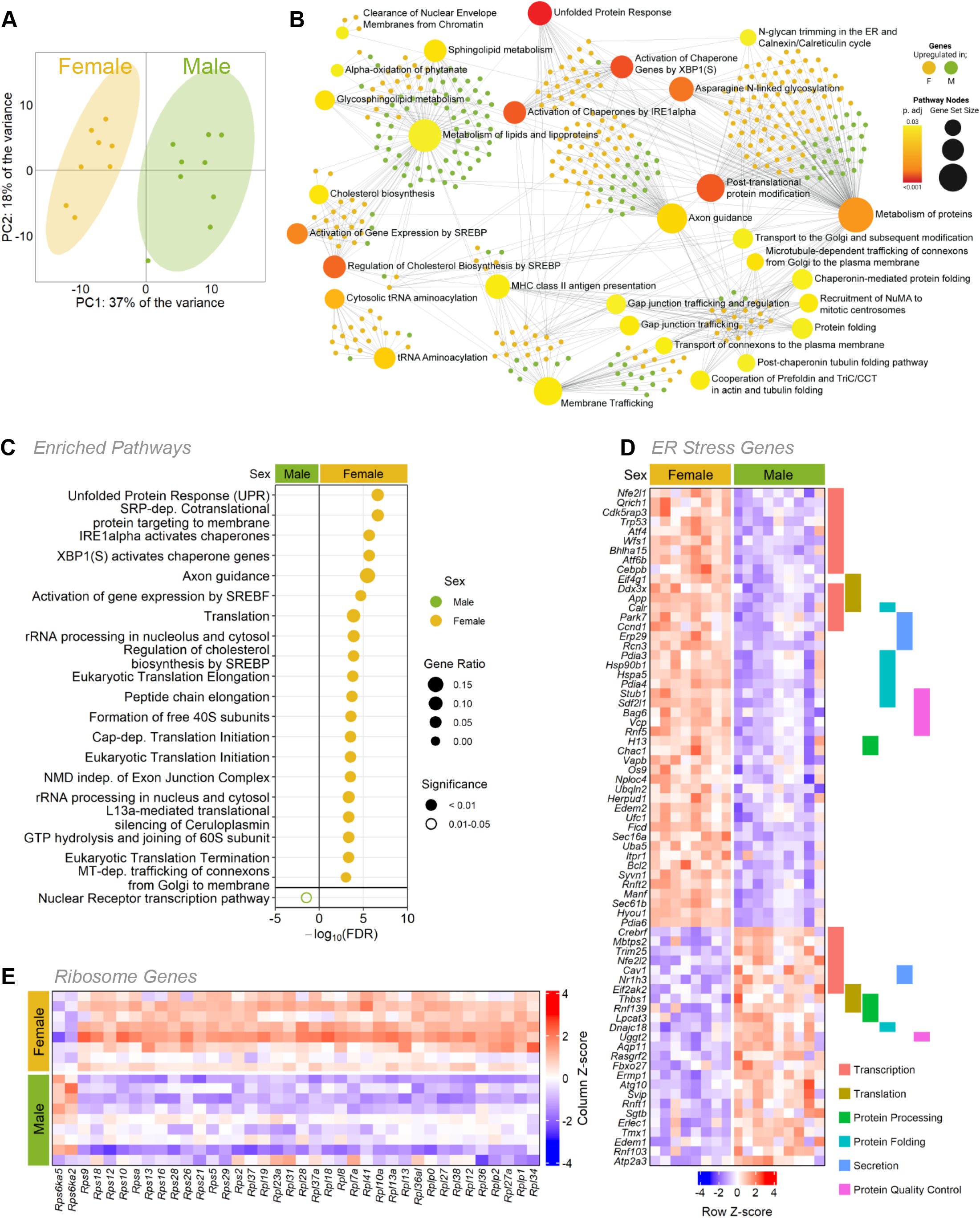
Sex-biased gene expression in mouse islet bulk RNAseq. (A) Principal component analysis (PCA) of RNAseq data from male and female mouse islets. (B) Over-representation analysis (ORA) of all significantly differentially expressed genes (*p*-adj < 0.01) from male and female mouse islets. Top 30 enriched KEGG pathways (large nodes; size = proportional to connections, darker red color = greater significance) and associated genes (small nodes; green = male enriched, yellow = female enriched). (C) Top significantly enriched Reactome pathways from the top 1000 significantly differentially expressed genes. (*p*-adj < 0.01) for males and females. Gene ratio is calculated as k/n, where k is the number of genes identified in each Reactome pathway, and n is the number of genes from the submitted gene list participating in any Reactome pathway. For complete Reactome pathway lists see Supplementary file 1. (D) All transcripts of differentially expressed genes under the gene ontology term “Cellular response to ER stress” (GO:0034976) and genes labeled by their role in transcription, translation, protein processing, protein folding, secretion and protein quality control. (E) All transcripts of differentially expressed ribosomal genes.

### 3.3. Female islets are more resilient to endoplasmic reticulum stress in mice

The burden of insulin production causes endoplasmic reticulum (ER) stress in β cells (68–70). ER stress is associated with an attenuation of mRNA translation (71), and, if ER stress is prolonged, can lead to cell death (72–74). Given that female islets exhibited higher expression of genes associated with protein synthesis, processing, and folding than males, and higher expression of genes associated with the UPR, which is activated in response to ER stress (75), we examined global protein synthesis rates in male and female islets under basal conditions and under ER stress. We incubated islets with O-propargyl-puromycin (OPP), which is incorporated into newly-translated proteins and can be ligated to a fluorophore. Using this technique, we monitored the accumulation of newly-synthesized islet proteins with single-cell resolution (Figure S6A). In basal culture conditions, male islet cells had significantly greater protein synthesis rates compared with female islet cells (Figure S6B). To investigate islet protein synthesis under ER stress in each sex, we treated islets with thapsigargin (Tg), a specific inhibitor of the sarcoplasmic/endoplasmic reticulum Ca^2+^-ATPase (SERCA) that induces ER stress and the UPR by lowering ER calcium levels (72,76). At 2 hours post-Tg treatment, protein synthesis was repressed in both male and female islet cells (Figure 3A, B; Figure S6C). At 24 hours post-Tg treatment, we found that protein synthesis was restored to basal levels in female islet cells, but not in male islet cells (Figure 3A, B; Figure S6C). Importantly, recovery from protein synthesis repression was significantly different in males and females (sex:treatment interaction p=0.0115). This suggests that while protein synthesis repression associated with ER stress was transient in female islets, this phenotype persisted for longer in male islets. Because insulin biosynthesis accounts for approximately half the total protein production in β cells (77), one potential explanation for the sex-specific recovery from protein synthesis repression is a sex difference in transcriptional changes to insulin. To test this, we quantified GFP levels in β cells isolated from male and female mice with GFP knocked into the endogenous mouse *Ins2* locus (*Ins2*^GFP/WT^)(51,78). While ER stress induced a significant reduction in *Ins2* gene activity, this response was equivalent between the sexes. This suggests *Ins2* transcriptional changes cannot fully explain the sex difference in recovery from protein synthesis repression during ER stress (Figure S7).

**Figure 3.**
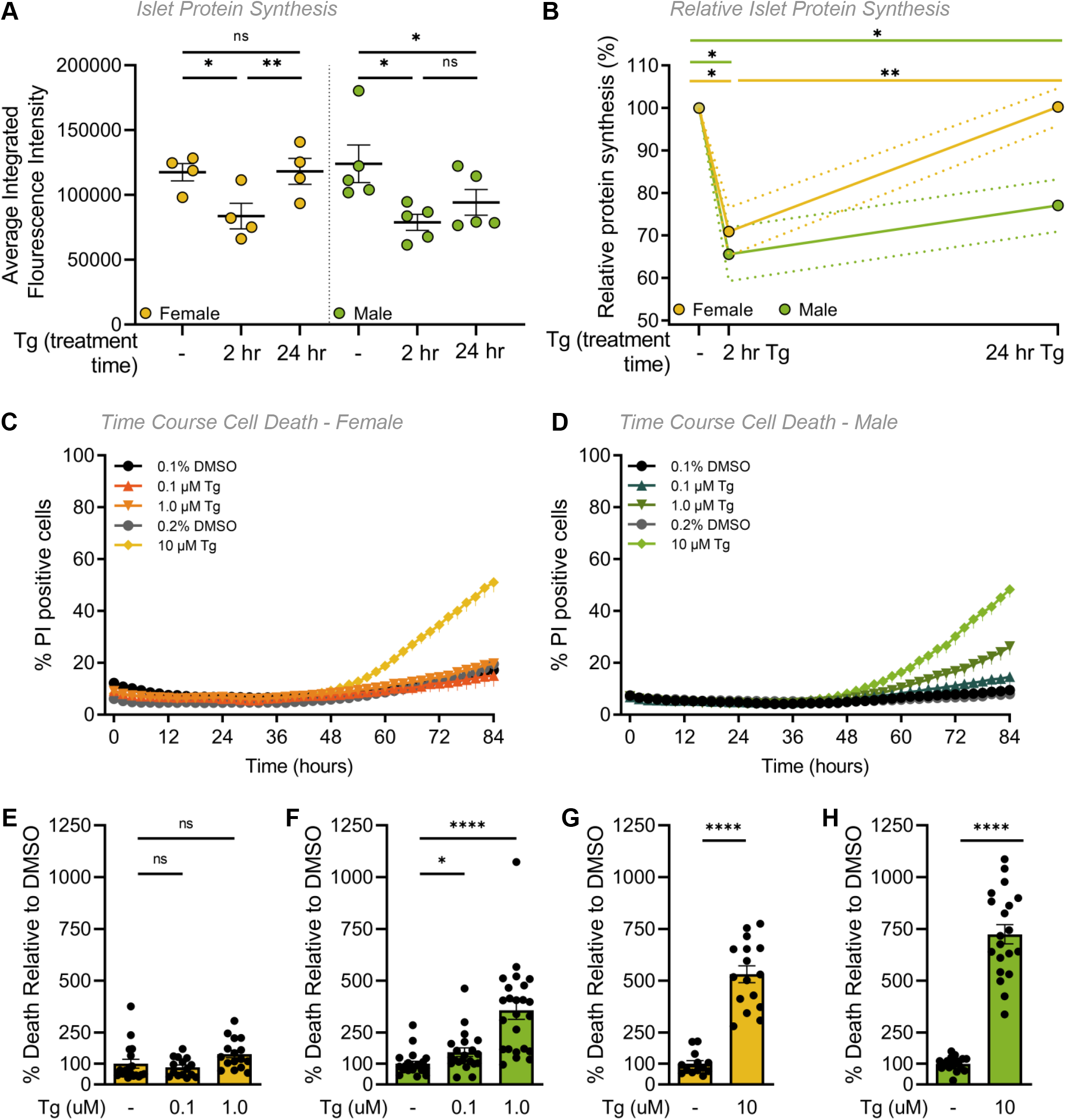
Sex differences in mouse islet ER stress-associated phenotypes. (A) Protein synthesis was quantified in dispersed islet cells from 20-week-old male and female B6 mice after treatment with 1 μM Tg for 2- or 24-hours. In female islet cells, protein synthesis was significantly lower after a 2-hour Tg treatment compared to control (p=0.0152; paired Student’s *t*-test) and significantly higher after a 24-hour Tg treatment compared to a 2-hour Tg treatment (p=0.0027; paired Student’s *t*-test). In male islet cells, protein synthesis was significantly lower after a 2- and 24-hour Tg treatment compared to control (p=0.0289 [0-2 hour] and p=0.0485 [0-24 hour]; paired Student’s *t*-test). (B) In both male and female islet cells protein synthesis was repressed after 2 hours. By 24 hours, protein synthesis repression was resolved in female, but not male islet cells. (C-H) Quantification of propidium iodide (PI) cell death assay of dispersed islets from 20-week-old male and female B6 mice treated with thapsigargin (0.1 μM, 1 μM or 10 μM Tg) or DMSO for 84 hours. n=4-5 mice, >1000 cells per group. Percentage (%) of PI positive cells was quantified as the number of PI-positive/Hoechst 33342-positive cells in female (C) and male (D) islet cells. Relative cell death at 84 hr in Tg treatments compared with DMSO treatment in females (E, G) and males (F, H). In female islet cells, cell death was significantly higher in 10 μM Tg compared to control (p<0.0001; unpaired Student’s *t*-test). In male islet cells, cell death was significantly higher in 0.1, 1.0 and 10 μM Tg compared to control (p=0.0230 [0.1 μM], p<0.0001 [1 μM] and p<0.0001 [10 μM]; unpaired Student’s *t*-test) (D). For E-H, at 84 hours the % of PI positive cells for each treatment was normalized to the DMSO control avg for each sex. * indicates p<0.05, ** indicates p<0.01, **** indicates p<0.0001; ns indicates not significant; error bars indicate SEM.

Given the prolonged protein synthesis repression in males following ER stress, we next quantified cell death, another ER stress-associated phenotype (75), in male and female islets. Using a kinetic cell death analysis, we observed clear sex differences in Tg-induced cell death at 0.1 μM and 1.0 μM Tg doses throughout the time course of the experiment (Figure 3C, D). After 84 hours of Tg treatment, no significant increase in female islet cell apoptosis was observed with either 0.1 μM or 1.0 μM Tg treatment compared with controls (Figure 3E). In contrast, cell death was significantly increased at both the 0.1 μM and the 1.0 μM doses of Tg in male islet cells compared with vehicle-only controls (Figure 3F). Importantly, our analysis shows the magnitude of Tg-induced cell death was larger in male islet cells compared with female islet cells (sex:treatment interaction p=0.0399 [0.1 μM], p=0.0007 [1.0 μM]). While one possible explanation for these data is that female islets are resistant to Tg-induced cell death, we found a significant increase in apoptosis in both female and male islet cells treated with 10 μM Tg (Figure 3G, H, sex:treatment interaction p=0.0996 [0.1 μM]). This suggests female islets were more resilient to mild ER stress caused by low-dose Tg than male islets.

To determine whether this increased ER stress resilience was caused by differential UPR signaling, we monitored levels of several protein markers of UPR activation including binding immunoglobulin protein (BiP), phosphorylated inositol-requiring enzyme 1 (pIRE1), phosphorylated eukaryotic initiation factor alpha (peIF2α), and C/EBP homologous protein (CHOP) (79,80) after treating male and female islets with 1 μM Tg for 24 hours. We found no sex difference in UPR protein markers between male and female islets without Tg treatment (Figure S8A-D) and observed a significant increase in levels of pIRE1α and CHOP in islets from both sexes and BiP in female islets after a 24-hour Tg treatment (Figure S8A-D). Lack of a sex difference in protein markers suggests UPR activation by Tg treatment was similar between male and female islets at 20 weeks of age. While this finding differs from other studies showing male-biased UPR activation (37), we reproduced the male-biased induction of BiP in islets isolated from 60-week-old male and female mice (Figure S8E-G), suggesting that age plays a role in the sex difference in UPR activation. Together, our data indicate that despite equivalent UPR activation in male and female islets treated with Tg, significant sex differences exist in ER stress-associated protein synthesis repression and cell death.

### 3.4. Female islets retain greater β cell function during ER stress in mice

We next examined glucose-stimulated insulin secretion in islets cultured under basal conditions and after Tg treatment (Figure 4A). In all conditions tested, high glucose significantly stimulated insulin secretion in both sexes (Figure S9A); however, we identified sex differences in how well islets sustained glucose-stimulated insulin secretion during longer Tg treatments (Figure 4B, C, Figure S9A). Female islets, in both low and high glucose, maintained robust insulin secretion during Tg treatment (Figure 4B). Specifically, we observed a significant increase in insulin secretion after short Tg treatment (0 and 2 hours post-Tg), with a return to basal secretion levels 4 hours post-Tg (Figure 4B). In contrast, male islets showed no significant increase in insulin secretion after short Tg treatment, and there was a significant drop in insulin secretion at 4 hours post-Tg treatment (Figure 4C). This suggests female islets sustained insulin secretion for a longer period than male islets during ER stress. Given that insulin content measurements showed insulin content significantly increased during the 4-hour Tg treatment in female islets, but not male islets (Figure 4D), our data suggest one reason female islets maintain insulin secretion during ER stress is by augmenting islet insulin content. Proinsulin secretion followed similar trends to what we observed with insulin secretion (Figure S9B), but Tg treatment reduced proinsulin content to a greater degree in male islets (Figure 4E). This suggests that in addition to females maintaining better insulin secretion during ER stress, they also show a larger increase in insulin content and a smaller decrease in proinsulin content.

**Figure 4.**
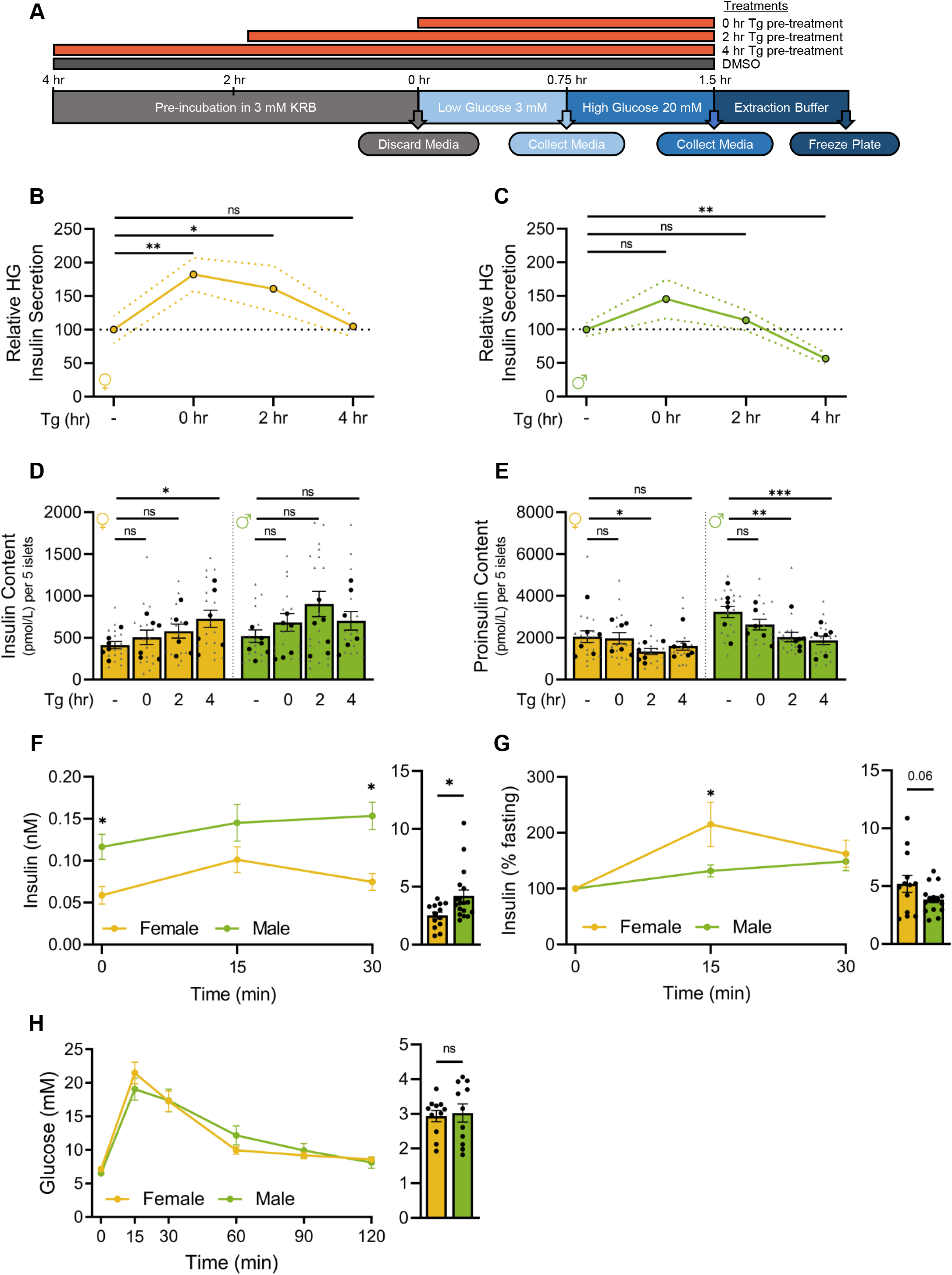
Sex differences in *ex vivo* and *in vivo* insulin secretion. (A) Experimental workflow of static glucose-stimulated insulin secretion. (B, C) Relative high glucose (20 mM; high glucose, HG) in treatments compared with DMSO in female (B) and male (C) islets. Female islet HG secretion was significantly higher compared with control after 0- and 2-hour Tg pre-treatments (p=0.0083 [0-hour] and p=0.0371 [2-hour]; Mann Whitney test). Male islet HG secretion was significantly lower compared with control after a 4-hour Tg pre-treatment (p=0.0013; Mann Whitney test). (D) Insulin content. Female islet insulin content was significantly higher compared with control after a 4-hour Tg pre-treatment (p=0.0269; Mann Whitney test). (E) Proinsulin content. Female islet proinsulin content was significantly lower compared with control after a 2-hour Tg pre-treatment (p=0.0437; Mann Whitney test). Male islet proinsulin content was significantly lower compared with control after 2- and 4-hour Tg pre-treatments (p=0.0014 [2-hour] and p=0.0005 [4-hour]; Mann Whitney test). (F-H) Physiology measurements after a 6-hour fast in 20-week-old male and female B6 mice. (F, G) Insulin levels from glucose-stimulated insulin secretion tests (F: nM, G: % basal insulin) following a single glucose injection (2 g glucose/kg body weight, i.p). Area under the curve (AUC) calculations (n=13 females, n=18 males). (F) Insulin levels were significantly higher in male mice at 0 minutes and 30 minutes post injection (p=0.0063 [0 minutes] and p=0.0009 [30 minutes]; Student’s *t*-test). AUC was significantly higher in males (p=0.0159; Student’s *t*-test). (G) Insulin levels (% baseline). Glucose-stimulated insulin secretion was significantly higher in female mice 15 minutes post injection (p=0.0279; Student’s *t*-test). (H) Glucose levels from glucose tolerance tests following a single glucose injection (2 g glucose/kg body weight). AUC calculations (n=11 females, n=11 males). For B-E, grey triangles indicate the concentration of insulin or proinsulin from five islets, black circles indicate the average values per mouse. For B, ## indicates p<0.01 and ### indicates p<0.001 for comparisons between treatments and DMSO in low glucose. For all other figures, * indicates p<0.05, ** indicates p<0.01, *** indicates p<0.001; ns indicates not significant; error bars indicate SEM.

To determine whether female islets show improved β cell function under ER stress in other contexts, we next monitored glucose-stimulated insulin secretion and glucose tolerance in mice at 20 weeks, an age where we show insulin sensitivity was equivalent between the sexes (Figure 4F-H; Figure S4). Despite higher fasting plasma insulin levels in males (Figure 4F), and similar glucose tolerance (Figure 4H), we found that the magnitude of glucose-stimulated insulin secretion was greater in females (Figure 4G). Given that ER stress exists even in normal physiological conditions due to the burden of insulin production (81), this adds further support to a model in which female β cells maintain better insulin production than male β cells under ER stress.

### 3.5. Sex differences in islet transcriptional and proteomic responses to ER stress in mice

To gain insight into the differential ER stress-associated phenotypes in male and female islets, we investigated global transcriptional changes after either a 6- or 12-hour Tg treatment in each sex. Principal component analysis and unsupervised clustering shows that islets clustered by sex, treatment, and treatment time (Figure 5A; Figure S10A). The majority of the variance was explained by treatment (Figure 5B), and pathway enrichment analysis confirms the UPR as the top upregulated pathway in Tg-treated male and female islets at both 6- and 12-hours after treatment (Figure S11A, B; Supplementary file 3). While some UPR-associated genes differentially regulated by Tg treatment were shared between the sexes (6-hour: 29/36, 12-hour: 25/31), biological sex explained a large proportion of variance in the gene expression response to ER stress. This suggests the transcriptional response to ER stress was not fully shared between the sexes. Indeed, after a 6-hour Tg treatment, 32.6% (2247/4655) of genes that were differentially expressed between DMSO and Tg were unique to one sex (881 to females, 1376 to males). After a 12-hour Tg treatment, 29% (2259/7785) were unique to one sex (1017 to males, 1242 to females).

**Figure 5.**
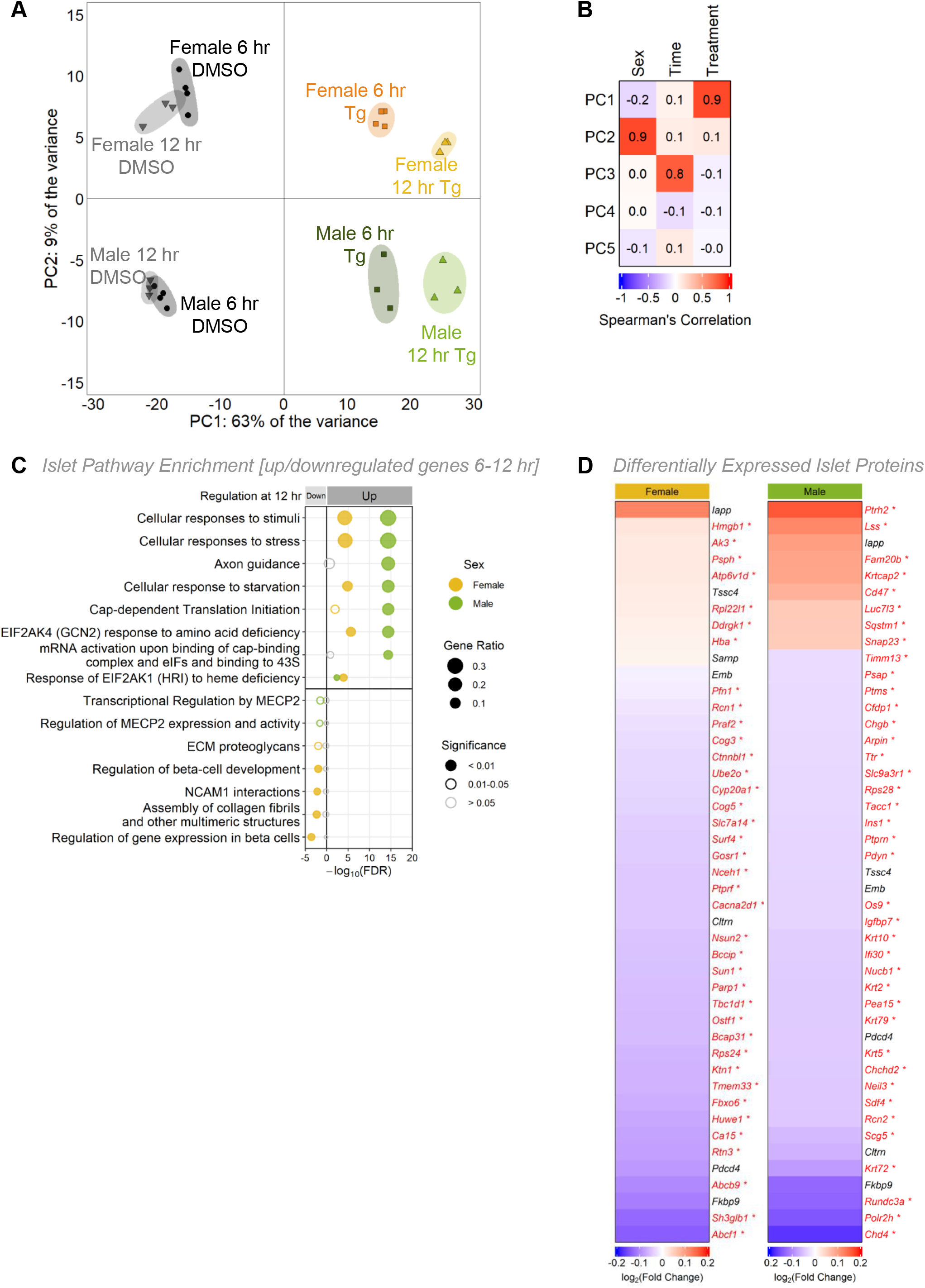
Sex-specific transcriptomic and proteomic profiles following ER stress in mouse islets. (A) Principal component analysis (PCA) of RNAseq data from male and female mouse islets treated with DMSO or 1 μM Tg for 6- or 12-hours. (B) Spearman correlation depicting the variance for the first 5 principal components. (C) Top significantly enriched Reactome pathways from the top 1000 significantly differentially expressed genes (*p*-adj<0.01) for females and males that were upregulated or downregulated between 6-12 hours of Tg treatment. Gene ratio is calculated as k/n, where k is the number of genes identified in each Reactome pathway, and n is the number of genes from the submitted gene list participating in any Reactome pathway. (D) Protein abundance from proteomics data of female and male mouse islets treated with DMSO or 1 μM Tg for 6 hours. Top 45 differentially expressed proteins are shown (p< 0.05).

To describe the transcriptional response of each sex to Tg treatment in more detail, we used a two-way ANOVA to identify genes that were upregulated, downregulated, or unchanged in male and female islets between 6- and 12-hours post-Tg (Supplementary file 4). By performing pathway enrichment analysis, we were able to determine which processes were shared, and which processes differed, between the sexes during Tg treatment. For example, we observed a significant increase in mRNA levels of genes corresponding to pathways such as cellular responses to stimuli, stress, and starvation in both male and female islets between 6- and 12-hour Tg treatments (Figure 5C; Supplementary file 4), suggesting Tg has similar effects on genes related to these pathways in both sexes. In contrast, there was a male-specific increase in mRNA levels of genes associated with translation during Tg treatment (Figure 5C; Supplementary file 4). In females, there was a decrease in mRNA levels of genes associated with β cell identity, such as *Pklr, Rfx6, Hnf4a, Slc2a2, Pdx1*, and *MafA* (Figure S12A), and in genes linked with regulation of gene expression in β cells (Figure 5C). Neither of these categories were altered between 6- and 12-hour Tg treatments in males (Figure 5C; Figure S12B). While this data suggests some aspects of the gene expression response to ER stress were shared between the sexes, we found that many genes corresponding to important cellular processes were differentially regulated during Tg treatment in only one sex.

Beyond sex-specific transcriptional changes following Tg treatment, ER stress also had a sex-specific effect on the islet proteome. Although the majority of proteins were downregulated by Tg treatment due to the generalized repression of protein synthesis under ER stress (Figure 5D), we identified 47 proteins (35 downregulated, 12 upregulated in Tg) that were differentially expressed in female islets and 82 proteins (72 downregulated, 10 upregulated in Tg) that were differentially expressed after Tg treatment in male islets (Supplementary table 1). Proteins downregulated only in females include proteins associated with GO term ‘endoplasmic reticulum to Golgi vesicle-mediated transport’ (GO:0006888) (BCAP31, COG5, COG3, GOSR1), whereas proteins downregulated only in males include proteins associated with GO terms ‘insulin secretion’ (GO:0030073) (PTPRN2, CLTRN, PTPRN) and ‘lysosome pathway’ (KEGG) (NPC2, CTSZ, LAMP2, PSAP, CLTA). Importantly, only seven differentially expressed proteins were in common between the sexes (Figure 5D). This suggests that as with our phenotypic and transcriptomic data, the proteomic response to Tg treatment was largely not shared between the sexes.

## 4. Discussion

Emerging evidence shows biological sex affects many aspects of β cell gene expression and function. Yet, many studies on β cells do not include both sexes, or fail to analyze male and female data separately. To address this gap in knowledge, the goal of our study was to provide detailed information on sex differences in islet and β cell gene expression and function in multiple contexts. In humans, we used a large scRNAseq dataset from ND and T2D donors to reveal significant male-female differences in the magnitude of gene expression changes, and in the identity of genes that were differentially regulated, between ND and T2D donors. This suggests β cell gene expression changes in T2D are not fully shared between the sexes. Given that our analysis shows β cells from female donors living with T2D maintain better insulin production than β cells from male donors living with T2D, our findings suggest female β cells are more resilient than male β cells in the context of T2D. In mice, our unbiased analysis of gene expression in islets from males and females with equivalent insulin sensitivity revealed sex differences in genes associated with the UPR under normal physiological conditions. This differential gene expression was significant, as female islets were more resilient to phenotypes caused by ER stress and UPR activation than male islets, showed sex-specific transcriptional and proteomic changes in this context, and maintained better insulin secretion. Collectively, these data suggest that in rodents, β cells from females are more resilient to ER stress. Considering the well-established links between ER stress and T2D (79,82–84), our data suggests a model in which female β cells maintain better function in T2D because they are more resilient to ER stress and UPR activation. While future studies are needed to test this model, and to assess the relative contribution of sex differences in β cells to the sex-biased risk of T2D, our findings highlight the importance of including both sexes in islet and β cell studies.

With respect to gene expression, including both sexes in our analysis of β cell gene expression in human ND and T2D allowed us to uncover genes that were differentially regulated in T2D in each sex. Because many of these genes may have been missed if the scRNAseq data was not analyzed by sex, our findings advance knowledge of β cell changes in T2D by identifying additional genes that are differentially regulated in this context. This knowledge adds to a growing number of studies that identify sex differences in β cell gene expression during aging in humans (12), and in mice fed either a normal (4,11) or a high fat diet (11). Further, given that our RNAseq on islets from male and female mice with equivalent insulin sensitivity identifies genes and biological pathways that align with previous studies on sex differences in murine β cell gene expression (4,11), our data suggests that sex differences in islet and β cell gene expression cannot be explained solely by a male-female difference in peripheral insulin resistance. Instead, there is likely a basal sex difference in β cell gene expression that forms the foundation for sex-specific transcriptional responses to perturbations such as ER stress and T2D. By generating large gene expression datasets from islet from male and female mice with equivalent peripheral insulin sensitivity and from islets subjected to pharmacological induction of ER stress, our studies provide a foundation of knowledge for future studies aimed at studying the causes and consequences of sex differences in islet ER stress responses and β cell function following UPR activation. This will provide deeper mechanistic insight into the sex-specific phenotypic effects reported in animal models of β cell dysfunction (35–39,85–88) and the sex-biased risk of diseases such as T2D that are associated with β cell dysfunction (12,22,89,90).

Beyond gene expression, our sex-based analysis of mouse islets allowed us to uncover male-female differences in ER stress-associated phenotypes (*e.g*. protein synthesis repression, cell death). While previous studies identify a sex difference in β cell loss in diabetic mouse models (37,39,91), and show that estrogen plays a protective role via estrogen receptor α (ERα) against ER stress to preserve β cell mass and prevent apoptosis in cell lines, mouse models, and human islets (39,91,92), we extend prior findings by showing that differences in ER stress-induced cell death were present in the context of equivalent insulin sensitivity between the sexes. This suggests sex differences in ER stress-associated phenotypes occur prior to male-female differences in peripheral insulin sensitivity. Indeed, islets isolated from males and females with equivalent sensitivity also show a sex difference in protein synthesis repression, a classical ER stress-associated phenotype (75). While estrogen affects insulin biosynthesis via ERα (93), future studies will need to determine whether estrogen also allows female islets to restore protein synthesis to basal levels faster than male islets following ER stress. We currently lack this knowledge, as most studies on UPR-mediated recovery from protein translation repression use single- and mixed-sex animal groups, or cultured cells (94–99).

Assessing whether the recovery of protein synthesis contributes to reduced cell death in female islets following ER stress will also be an important task for future studies, as prior studies suggest the inability to recover from protein synthesis repression increases ER-stress induced apoptosis (94). Ideally, this type of study would also monitor the activity of pathways known to regulate protein synthesis repression during ER stress. For example, while we did not detect any changes in levels of phosphorylated eIF2α (also known as Eif2s1), which is known to mediate UPR-induced protein synthesis repression (75), our chosen timepoints did not overlap with the rapid changes in phospho-eIF2α following ER stress published in other studies (100,101). A more detailed time course will therefore be necessary to assess p-eIF2α levels during ER stress in both sexes, and to test a role for phospho-eIF2α in mediating differences in protein synthesis repression. Ultimately, a better understanding of sex differences in ER stress-associated phenotypes in β cells will provide a mechanistic explanation for the strongly male-biased onset of diabetes-like phenotypes in mouse models of β cell ER stress (*e.g*. Akita, KINGS, Munich mice) (37,38,85). Given the known relationship between ER stress, β cell death, and T2D, studies on the male-female difference in β cell ER stress-associated phenotypes may also advance our understanding of the male-biased risk of developing T2D in some population groups.

A further benefit of additional studies on the sex difference in β cell ER stress responses will be to identify mechanisms that support β cell insulin production. In rodents, we found that female islets maintained high glucose-stimulated insulin secretion and increased insulin content following ER stress, whereas male islets showed significant repression of high glucose-stimulated insulin secretion under the same conditions. In humans, while a study using a mixed-sex group of T2D donors shows β cells experience ER stress associated with β cell dysfunction (102), we found that changes to β cell insulin secretion in T2D were not the same between the sexes. Specifically, the magnitude of the reduction in insulin release by β cells from female donors living with T2D was smaller than in β cells from male donors living with T2D. Together with our data from rodents, this suggests female β cells maintain enhanced insulin production and/or secretion in multiple contexts, and the increased β cell function cannot be solely attributed to a sex difference in peripheral insulin sensitivity.

Clues into potential ways that female β cells maintain improved insulin production and secretion emerge from our examination of the transcriptional response to ER stress in each sex. Our data shows that Tg treatment induces gene expression changes characteristic of ER stress (103), and revealed similar biological pathways that were upregulated in T2D donors. Furthermore, we identified significant differences between male and female islets in the transcriptional response to ER stress over time. One notable finding was that a greater number of β cell identity genes were downregulated between 6- and 12-hour Tg treatments in females, but not in males. Because most studies on the relationship between β cell identity and function used a mixed-sex pool of islets and β cells (68,104,105), more studies will be needed to test whether there are sex-specific changes to β cell identity during ER stress, and to determine the functional consequences of this sex-specific effect. Ultimately, a better understanding of changes to β cell gene expression and function in males and females will suggest effective ways to reverse disease-associated changes to this important cell type in each sex, improving equity in health outcomes (106).

### 4.1. Conclusions

Our study reports significant sex differences in islet and β cell gene expression and stress responses in both humans and mice. These differences likely contribute to sex differences in β cell resilience, allowing female β cells to maintain better insulin production across multiple contexts. This knowledge forms a foundation for future studies aimed at understanding how sex differences β cell function affect physiology and the pathophysiology of diseases such as T2D.

## Funding

This study was supported by operating grants to E.J.R. from the Michael Smith Foundation for Health Research (16876), Canadian Institutes of Health Research (GS4-171365), the Canadian Foundation for Innovation (JELF-34879), and to J.D.J. (PJT-152999) from the Canadian Institutes of Health Research, and core support from the JDRF Centre of Excellence at UBC (3-COE-2022-1103-M-B). J.D.J. was funded by a Diabetes Investigator Award from Diabetes Canada.

## Acknowledgements

We thank members of the Rideout and Johnson lab for valuable feedback. We thank our animal care staff for supporting our animal husbandry, UBC Biomedical Research Center for performing RNA sequencing, and UVic Genome BC Proteomics Center for performing proteomics. We acknowledge that our research takes place on the traditional, ancestral, and unceded territory of the Musqueam people; a privilege for which we are grateful.

## Conflict of interest

The authors declare no competing interest.

## Data Availability

Details of all statistical tests and *p*-values are provided in Supplementary file 5. All raw data generated in this study are available in Supplementary file 6. RNAseq data is available in Supplementary file 7 and Supplementary file 8.

## SUPPLEMENTAL FIGURE LEGENDS

**Figure S1.**
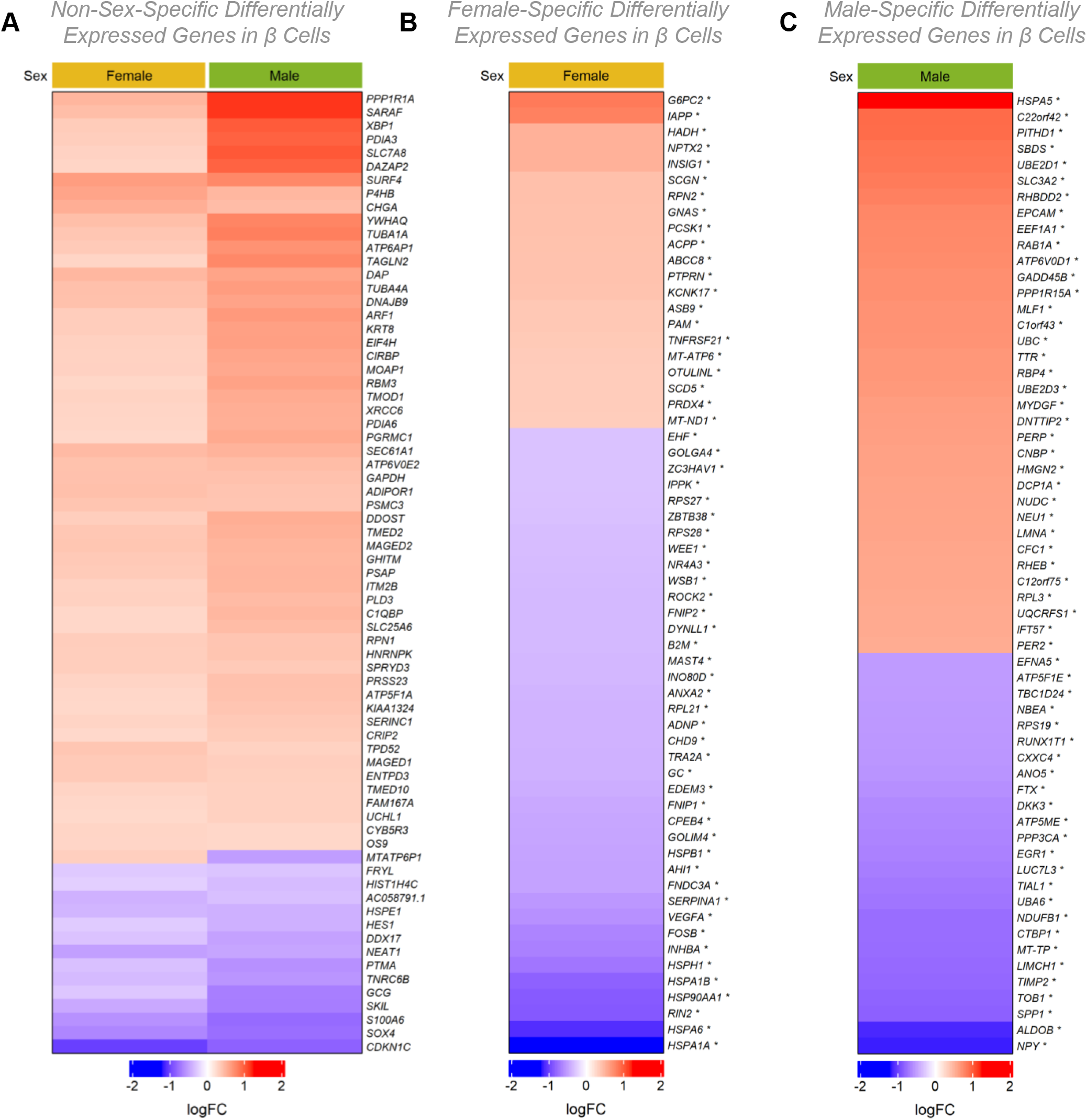
Sex-specific and non-sex-specific differentially expressed genes in T2D. scRNAseq data from male and female human β cells. (A-C) Top 60 significantly differentially expressed genes (*p*-adj < 0.05). Non-sex-specific (A), female-specific (B), or male-specific (C). For complete gene lists see Supplementary file 1.

**Figure S2.**
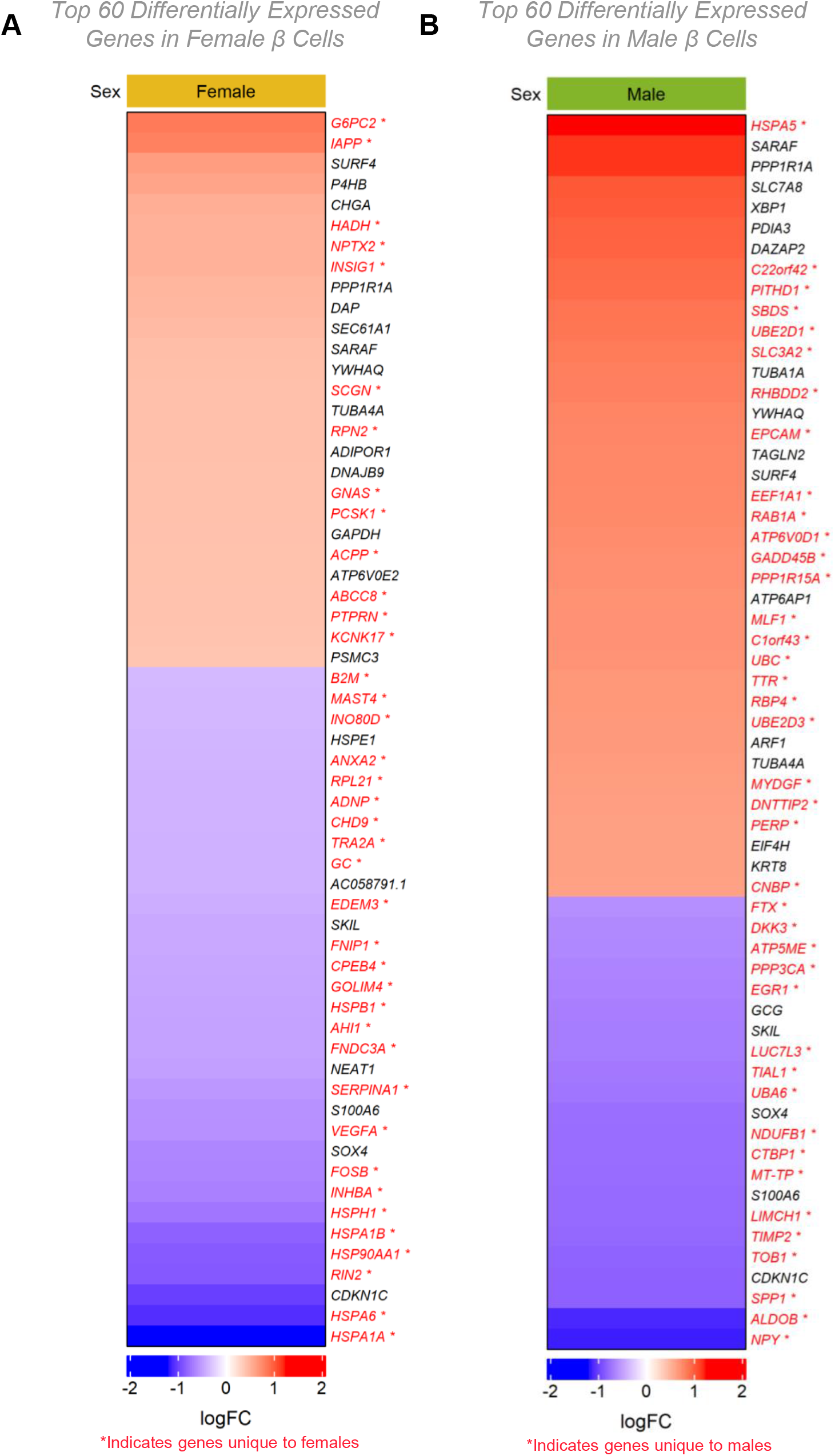
Gene expression changes in T2D. scRNAseq data from male and female human β cells. (A, B) Top 60 differentially expressed genes (*p*-adj < 0.05) in females (A) and males (B). Sex-specific genes are indicated in red text. For complete gene lists see Supplementary file 1.

**Figure S3.**
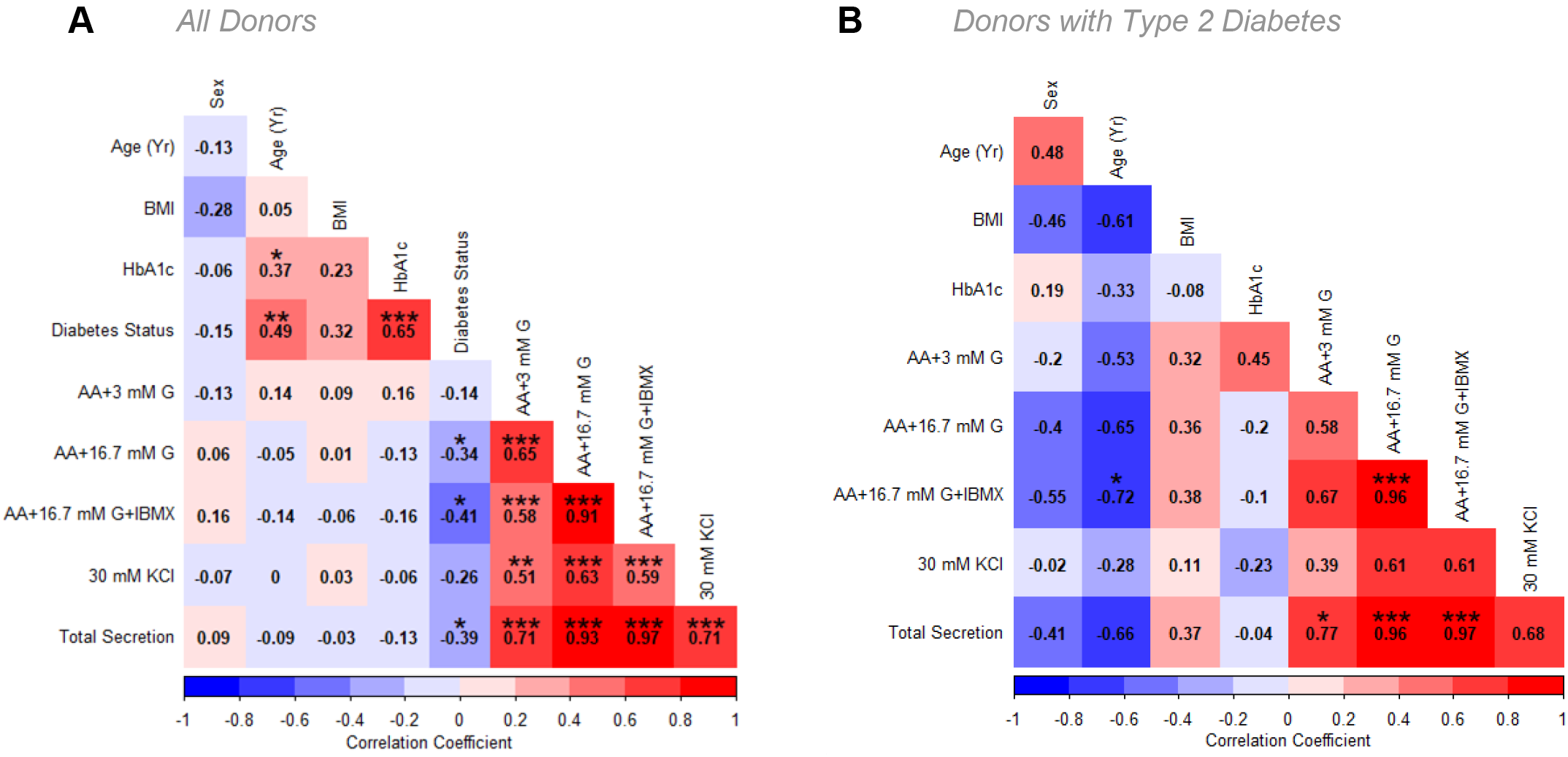
Correlations between donor attributes and insulin secretion. Pearson correlation of all human donors (A), or all donors with Type 2 Diabetes. Significant correlations are denoted with a star (*). For donor metadata see Supplementary file 6.

**Figure S4.**
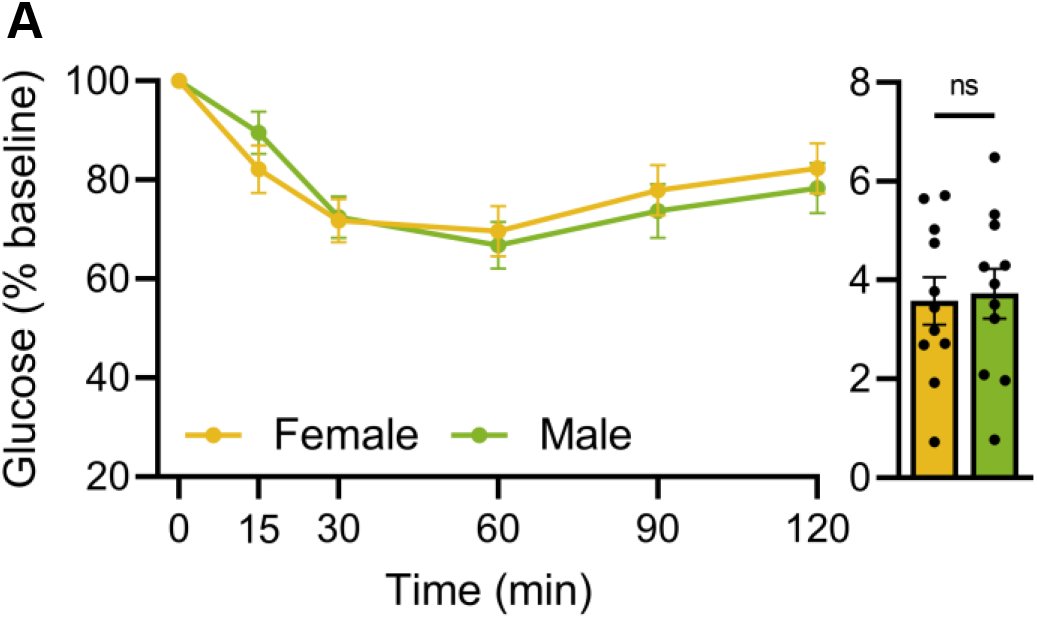
Equivalent insulin sensitivity in male and female mice. (A) Insulin tolerance test (ITT). 20-week-old female and male B6 mice were fasted for 6 hours. Glucose levels (% baseline) from insulin tolerance tests (ITT) following a single insulin injection (0.75U insulin/kg body weight). AUC calculations (n=11 females, n=11 males). ns indicates not significant; error bars indicate SEM.

**Figure S5.**
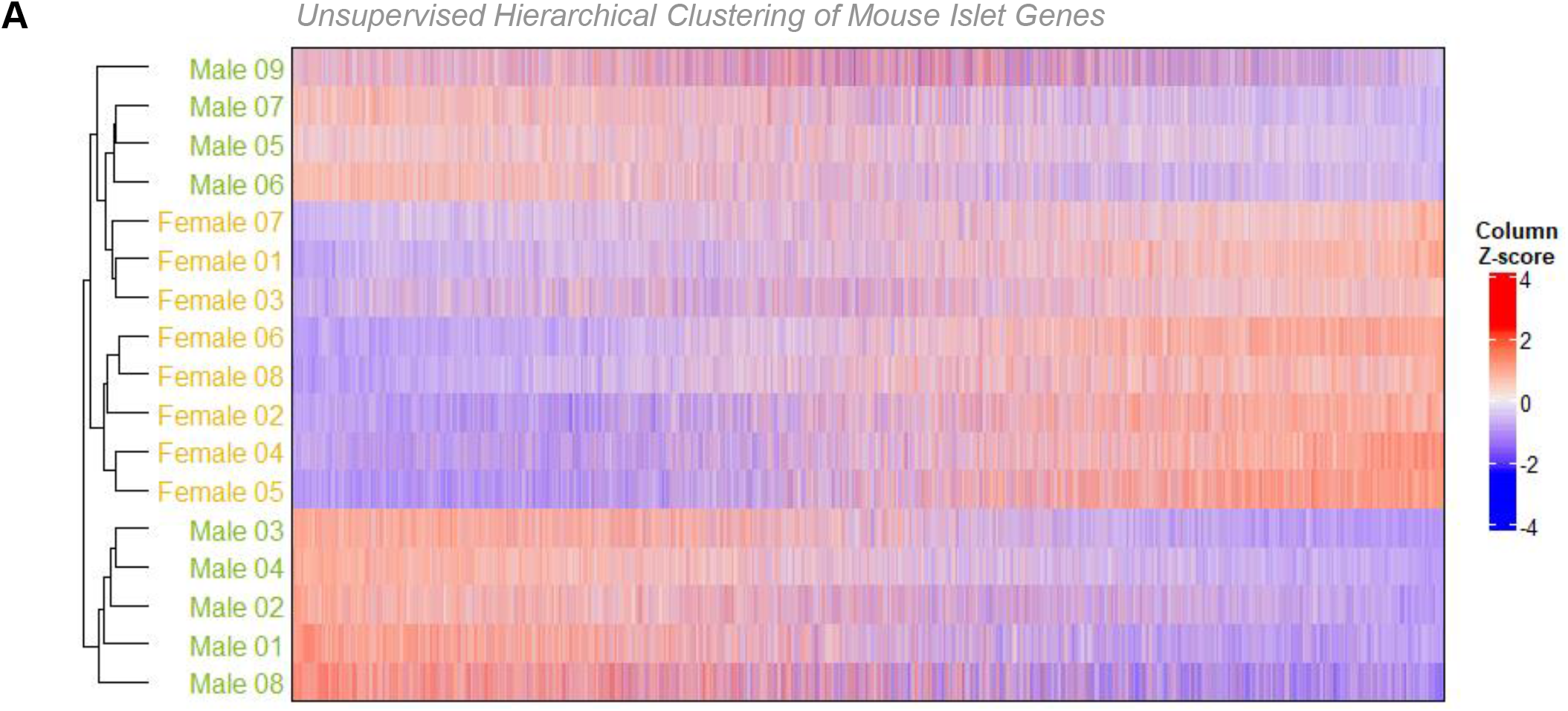
Mouse islet gene expression clusters by sex. (A) Unsupervised hierarchical clustering of RNAseq data from female and male mouse islets. Sorting was based on all genes where the total count was >10 across all samples.

**Figure S6.**
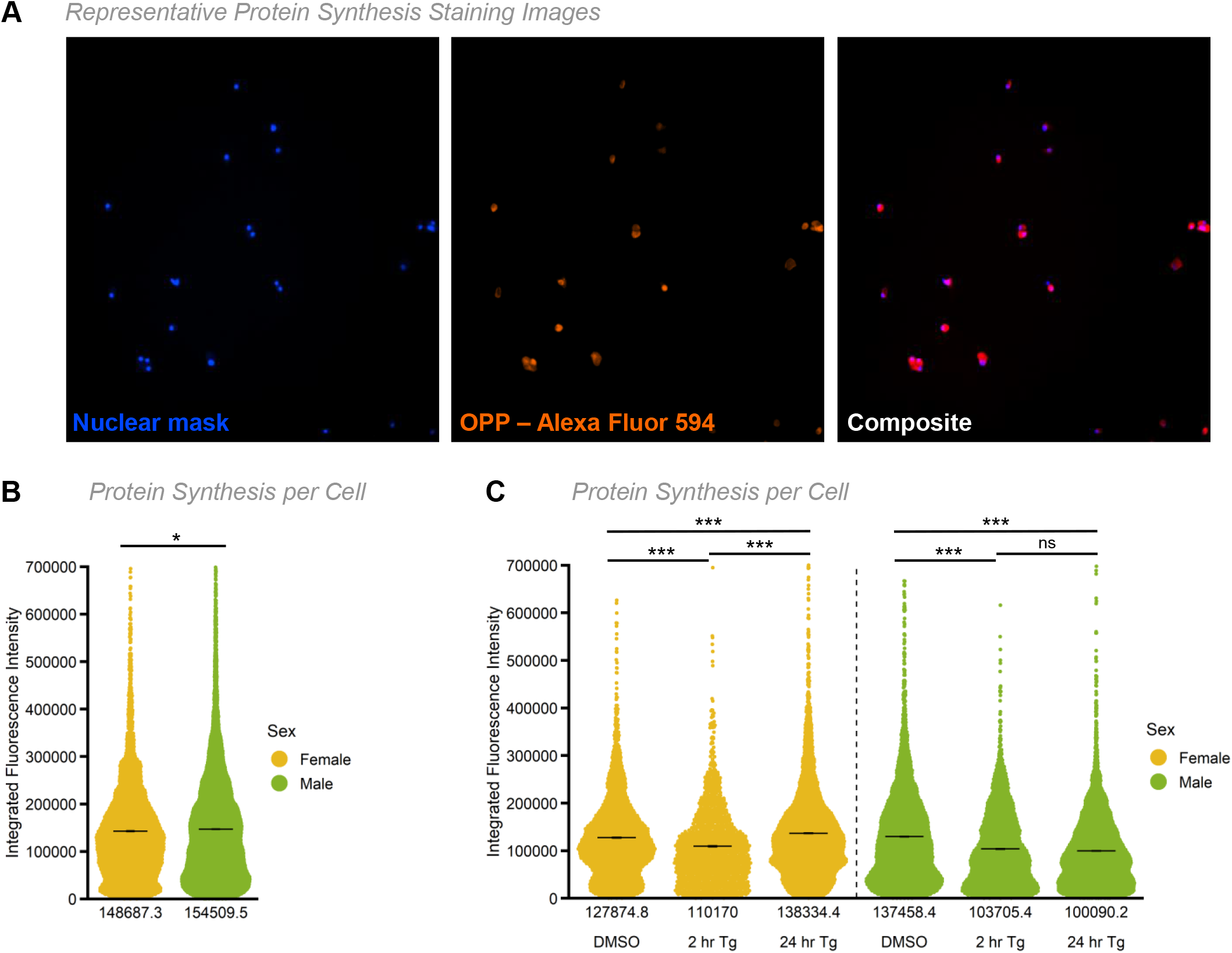
ER stress-induced protein synthesis repression persists in male mouse islet cells. (A) Representative images of dispersed islets stained with nuclear mask and OPP labeled with Alexa Fluor 594. (B, C) Integrated staining intensity of Alexa Fluor 594 in nuclear mask positive islet cells in control media (B, FBS+) or after treatment with DMSO control or 1 μM Tg for 2- or 24-hours (C, FBS-). Protein synthesis is displayed on a per cell basis from data shown in Figure 3. n=4-5 mice, >1000 cells per group. Mean values are indicated under each group. (B) Protein synthesis was significantly higher in male islet cells than female islet cells in control media, 3.9% (p=0.027; unpaired Student’s *t*-test). (C) In female islet cells, protein synthesis was significantly repressed from control-2 hour treatments (p=<0.0001; unpaired Student’s *t*-test) and significantly increased from both control-24 hour treatments and 2-24 hour treatments (p=<0.0001; unpaired Student’s *t*-test). In male islet cells, protein synthesis was significantly repressed from control-2 hour treatments (p=<0.0001; unpaired Student’s *t*-test) and control-24 hour treatments (p=<0.0001; unpaired Student’s *t*-test); however, was not significantly different between 2-24 hour treatments (p=0.07; unpaired Student’s *t*-test). * indicates p<0.05, *** indicates p<0.001; ns indicates not significant; error bars indicate SEM.

**Figure S7.**
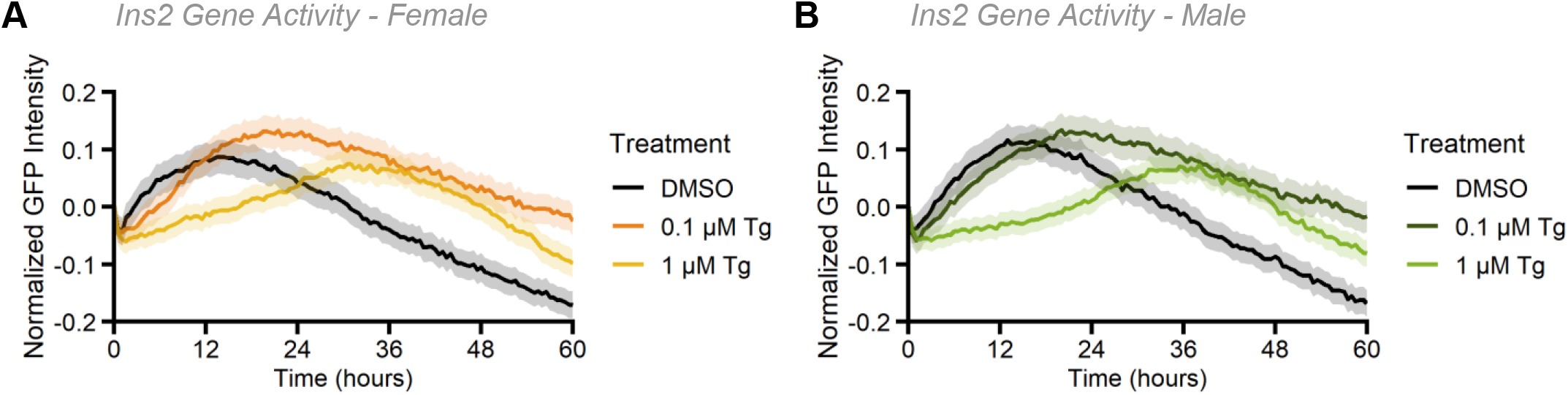
*Ins2* gene activity is repressed by ER stress induction. *Ins2* gene activity in β cells from 20-week-old male and female B6 mice treated with Tg (0.1 μM or 1 μM Tg) or DMSO for 60 hours (n=6 mice per sex, > 1000 cells per group). (A, B) Average change in fluorescence intensity from all GFP expressing female (A) and male (B) β cells over time. Data was normalized to the first 2 hours to examine relative change in *Ins2* gene activity. (C, D) Density plot of *Ins2*^GFP/WT^ β cell GFP fluorescence intensity, log transformed. Data is shown for each run for females (C) and males (D). (E-H) Average change in high (E, F) and low (G,H) GFP *Ins2*^GFP/WT^ β cells fluorescence over time from females (E, G) and males (F, H). Data was normalized to the first two hours to examine relative change in *Ins2* gene activity.

**Figure S8.**
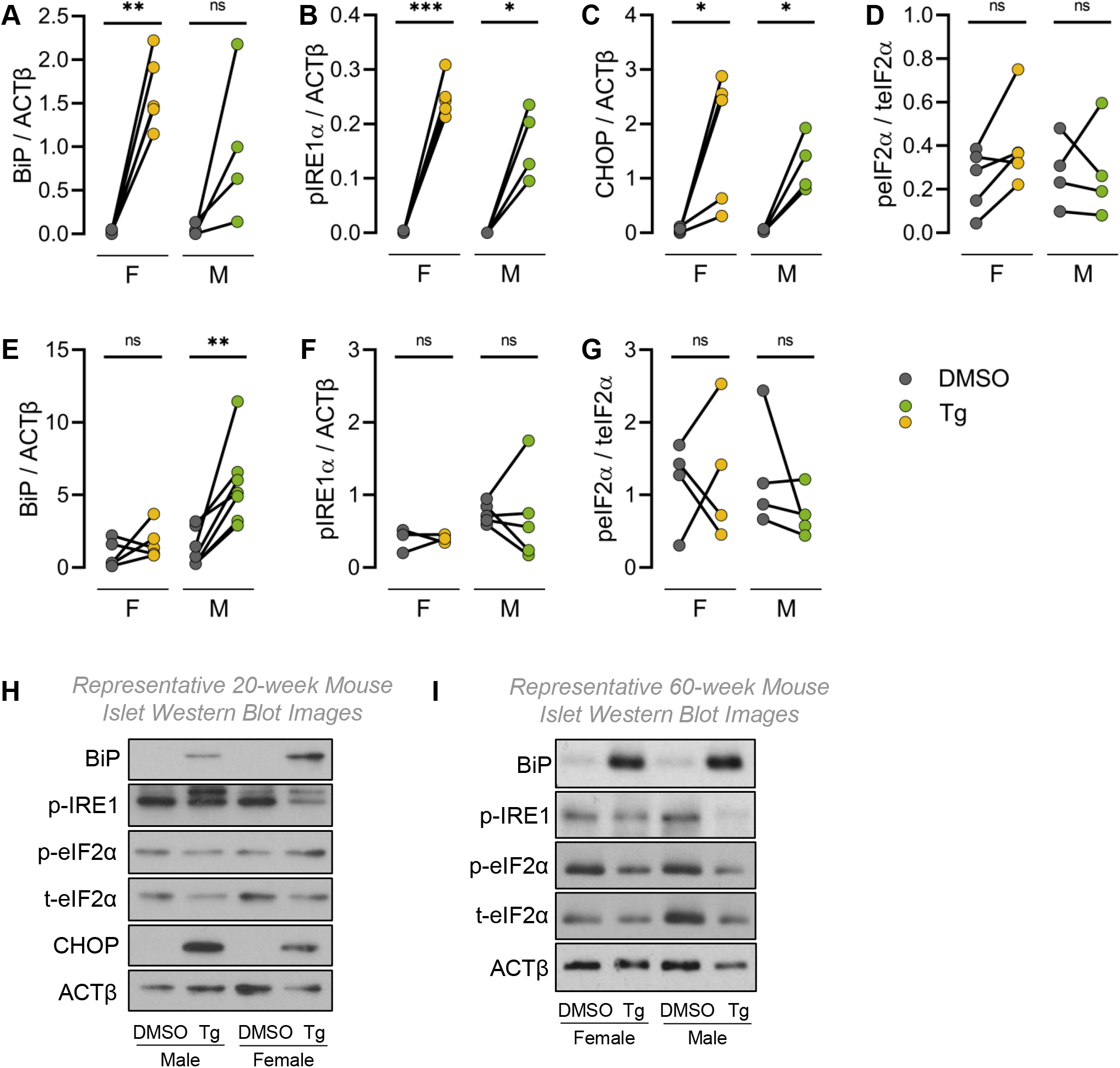
Representative western blot images of UPR protein markers. (A-D) Levels of ER stress proteins were quantified in isolated islets from 20-week-old male and female B6 mice cultured in DMSO or 1 μM Tg for 24 hours. (A) BiP levels were significantly upregulated in female Tg vs DMSO (p=0.0011; paired Student’s *t*-test) but not male Tg vs DMSO (p=0.1187; paired Student’s *t*-test). (B) pIRE1α levels were significantly upregulated in female Tg vs DMSO (p=0.0001; paired Student’s *t*-test) and in male Tg vs DMSO (p=0.0148; paired Student’s *t*-test). (C) CHOP levels were significantly upregulated in female Tg vs DMSO (p=0.0333; paired Student’s *t*-test) and in male Tg vs DMSO (p=0.0164; paired Student’s *t*-test). (D) p-eIF2α levels were not significantly upregulated in either sex. (E-G) Levels of ER stress proteins were quantified in isolated islets from 60-week-old male and female B6 mice cultured in DMSO or 1 μM Tg for 24 hours. (E) BiP levels were significantly upregulated in male Tg vs DMSO (p=0.0048; paired Student’s */*-test) but not in female Tg vs DMSO (p=0.3319; paired Student’s */*-test). (F) p-IRE1α levels were not significantly upregulated in either sex (p=0.9257 [female] and p=0.8273 [male]; paired Student’s *t*-test). (G) p-eIF2α levels were not significantly upregulated in either sex (p=0.8451 [female] and p=0.3076 [male]; paired Student’s *t*-test). (H) Representative western blot images of 20-week Tg treated mouse islets. (I) Representative western blot images of 60-week Tg treated mouse islets. * indicates p<0.05, ** indicates p<0.01; ns indicates not significant.

**Figure S9.**
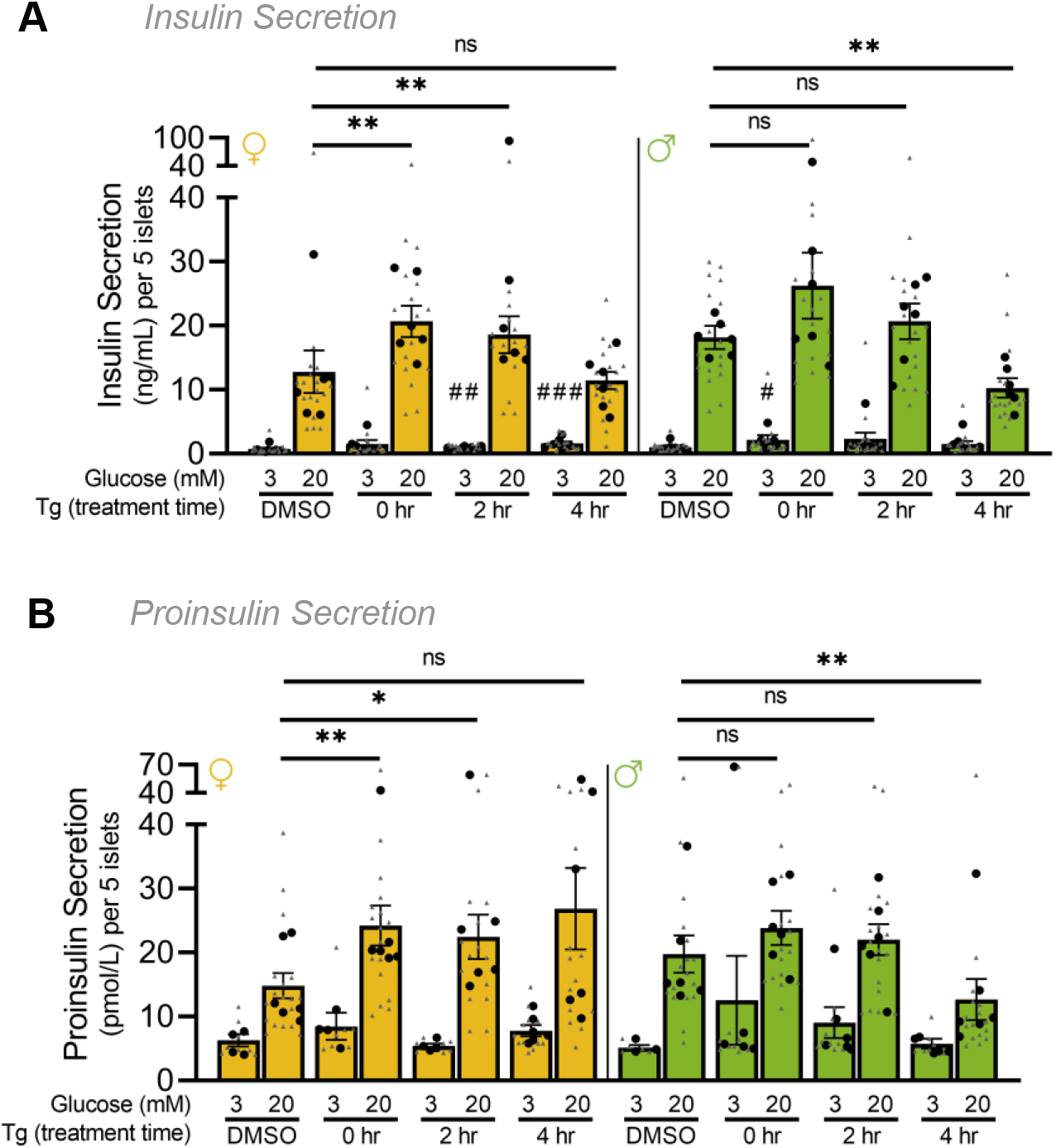
Female mouse islets retain greater insulin secretion during ER stress. (A) Insulin secretion at basal (3 mM; low glucose, LG) and stimulatory (20 mM; high glucose, HG) glucose. Female islet LG secretion was significantly higher compared with control after 2- and 4-hour Tg pre-treatments (p=0.0047 [2-hour] and p=0.0003 [4-hour]; Mann Whitney test). Female islet HG secretion was significantly higher compared with control after 0- and 2-hour Tg pre-treatments (p=0.0012 [0-hour] and p=0.0061 [2-hour]; Mann Whitney test). Male islet LG secretion was significantly higher compared with control after a 0-hour Tg pre-treatment (p=0.0371; Mann Whitney test). Male islet HG secretion was significantly lower compared with control after a 4-hour Tg pre-treatment (p=0.0012; Mann Whitney test). (B) Proinsulin secretion at basal (3 mM) and stimulatory (20 mM) glucose. Female islet HG secretion was significantly higher compared with control after 0- and 2-hour Tg pre-treatments (p=0.0075 [0hour] and p=0.0437 [2-hour],; Mann Whitney test). Male islet HG secretion was significantly lower compared with control after a 4-hour Tg pre-treatment (p=0.0025; Mann Whitney test). * indicates p<0.05, ** indicates p<0.01; ns indicates not significant; error bars indicate SEM.

**Figure S10.**
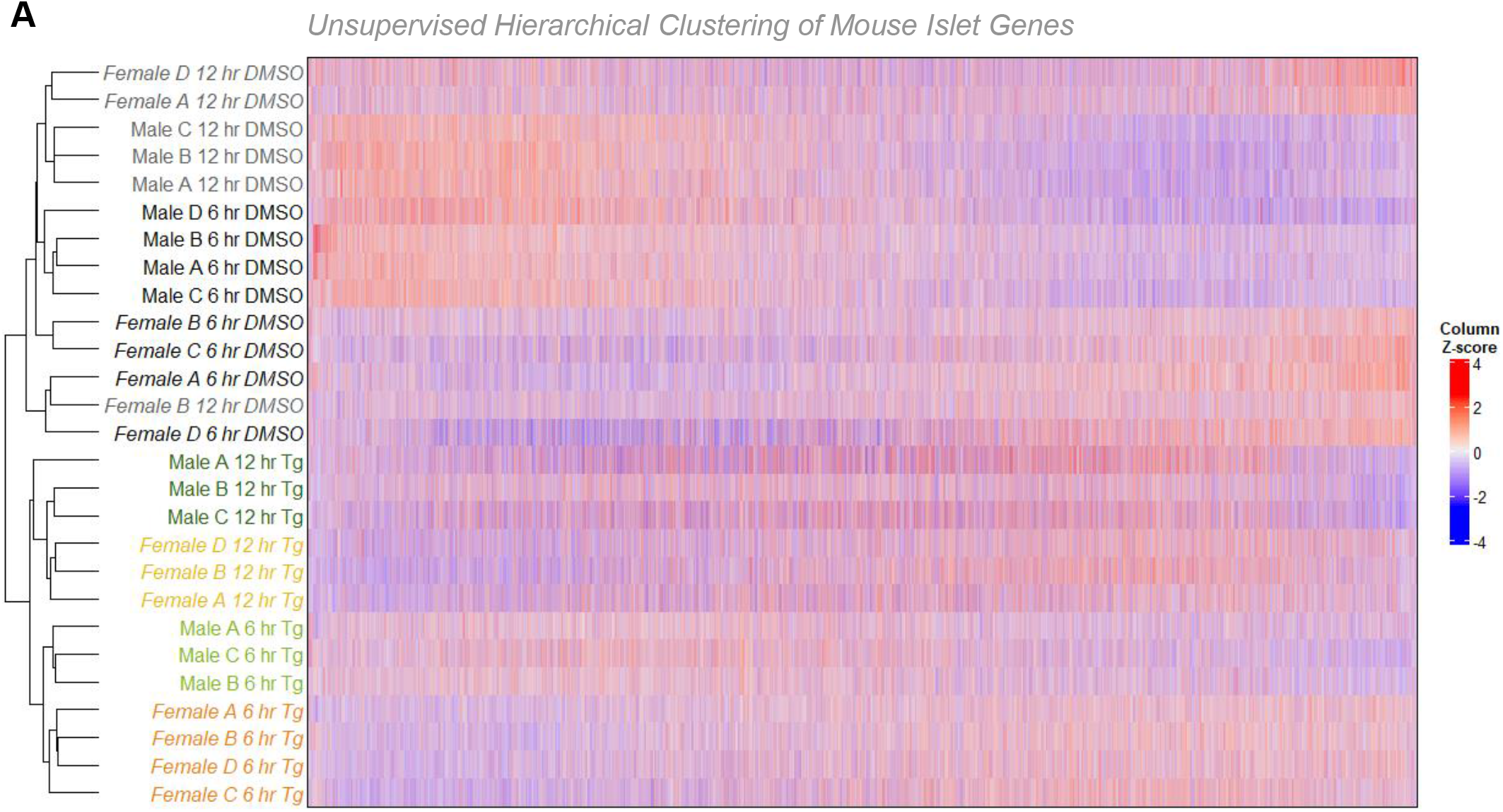
Mouse islet gene expression clusters by sex, treatment and time. (A) Unsupervised hierarchical clustering of RNAseq data from female and male DMSO or Tg treated mouse islets. Sorting was based on all genes where the total count >10 across all samples.

**Figure S11.**
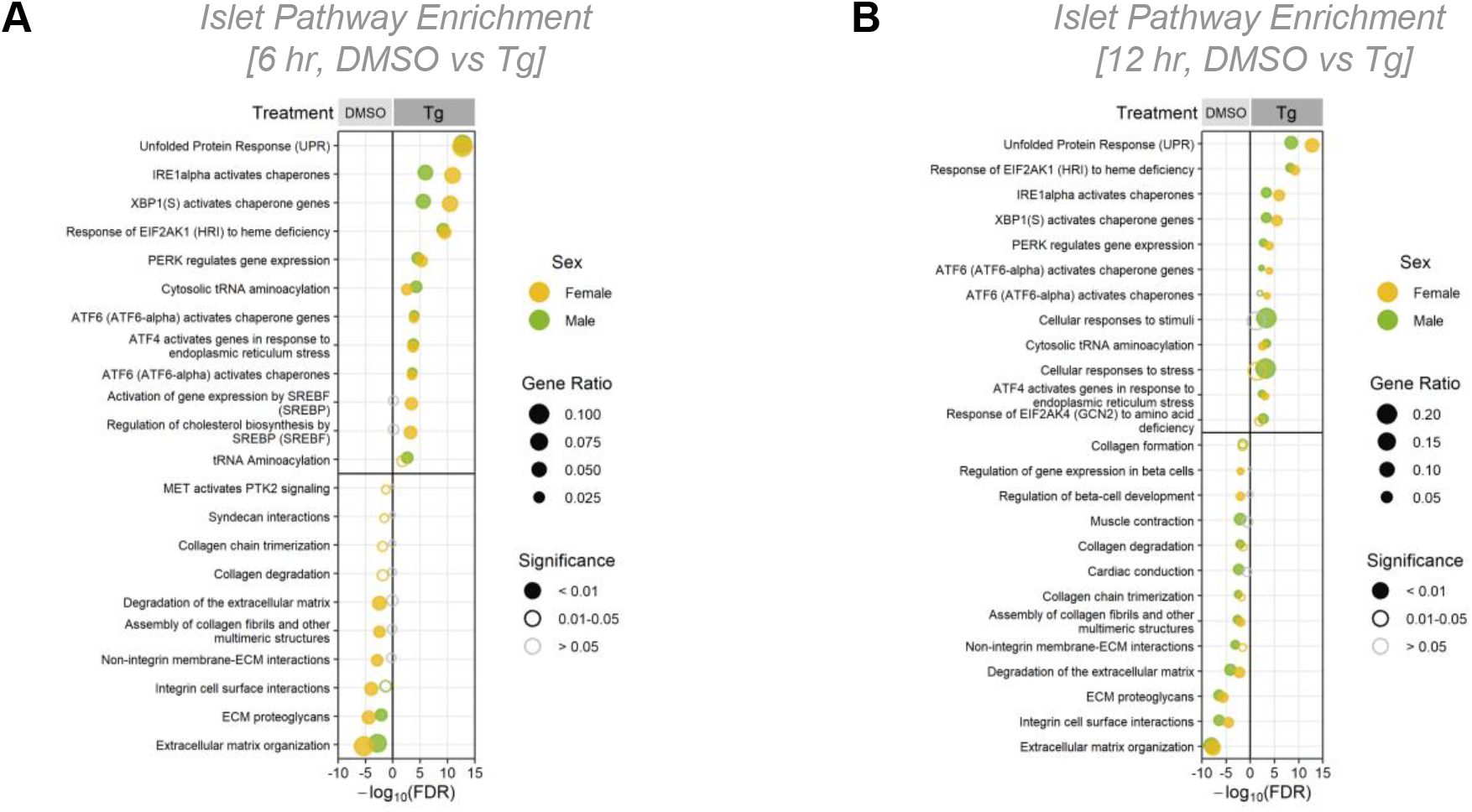
Female and Male mouse islets are enriched in similar pathways following 6- and 12-hour Tg treatments. (A, B) Most significantly enriched Reactome pathways from the top 1000 significantly differentially expressed genes. (*p*-adj < 0.01) for females and males between DMSO vs Tg after 6 hours (A) or 12 hours (B) of Tg treatment. Gene ratio is calculated as k/n, where k is the number of genes identified in each Reactome pathway, and n is the number of genes from the submitted gene list participating in any Reactome pathway.

**Figure S12.**
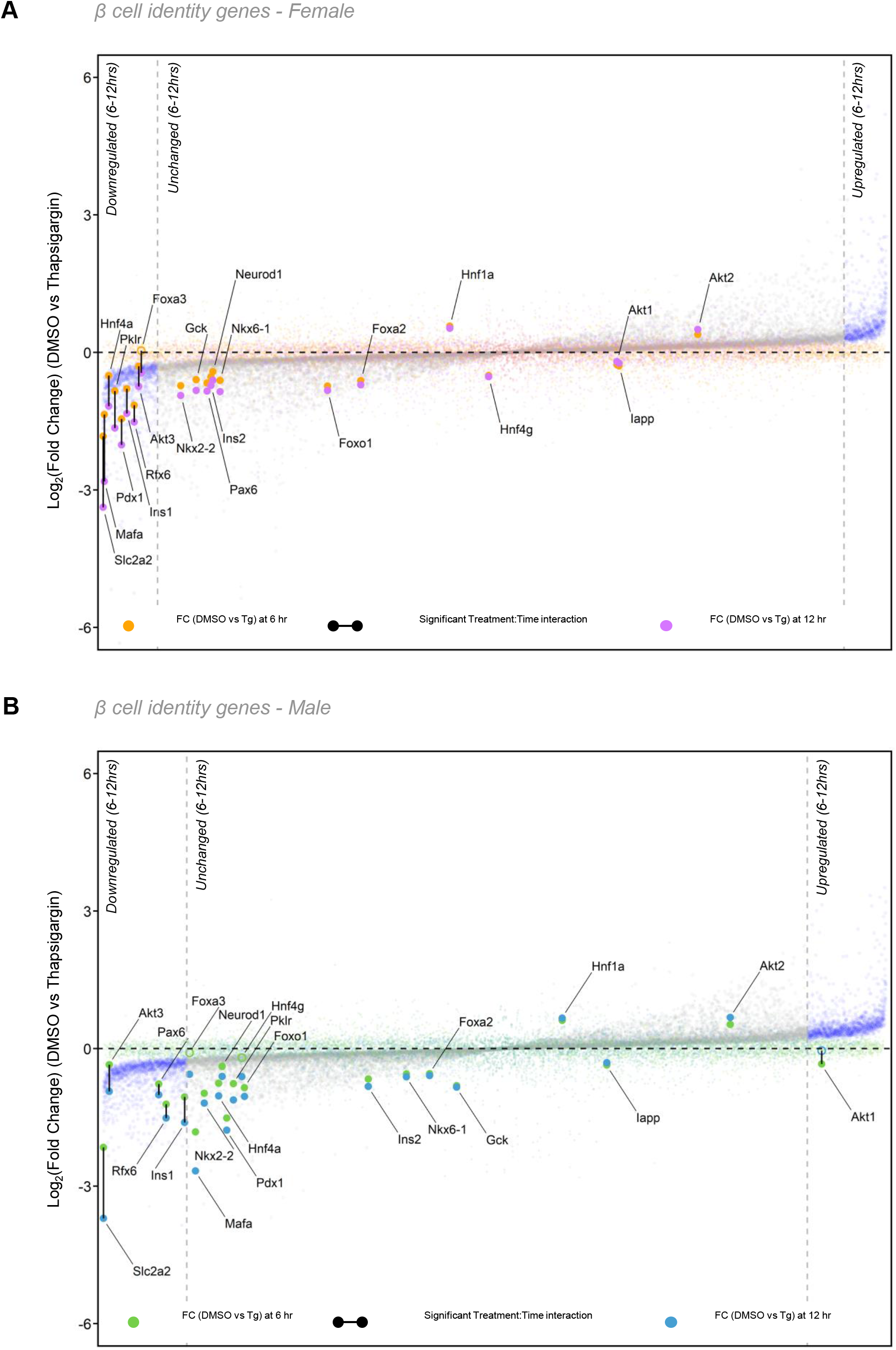
A greater number of β cell identity genes are downregulated between 6- and 12-hour Tg treatment times in female mouse islets. (A, B) Treatment:Time interaction plots of female islet (A) and male islet (B) β cell identity genes in Reactome pathway “Regulation of gene expression in β cells”. The fold change (FC) for DMSO vs Tg was calculated for each sex and time point (Female 6-hour, Female 12-hour, Male 6-hour, Male 12-hour). The change in FC values (12-hour FC – 6-hour FC) were plotted according to *p*-adj values. In females, FC values between 6- and 12-hours are represented by orange and purple dots, respectively. In males, FC values at 6- and 12-hours are represented by green and blue dots, respectively. A solid black line connecting the dots indicates genes with a significant treatment:time interaction.

## Notes

https://hpap.pmacs.upenn.edu/

### Competing Interest Statement

The authors have declared no competing interest.

### Summary of Updates

Updated figures, wording and layout of the manuscript.

